# Intraspecific variation in the duration of epigenetic inheritance

**DOI:** 10.1101/2025.06.04.657799

**Authors:** François Frejacques, Marie Saglio, Mohammed Aljohani, Christian Froekjaer-Jensen, Lise Frézal, Marie-Anne Félix

**Affiliations:** IBENS, Department of Biology, Ecole Normale Supérieure, CNRS, Inserm, PSL Research University, Paris, France; Bioscience Program, Biological and Environmental Science and Engineering (BESE), King Abdullah University of Science and Technology (KAUST), Thuwal, 23955-6900, Kingdom of Saudi Arabia; Department of Developmental and Stem Cell Biology, Institut Pasteur, Paris, France; Institut Pasteur, Université Paris Cité, Unité des Bactéries pathogènes entériques, Paris, France; Department of Genome Sciences, University of Washington, Seattle, WA 98195, USA

**Keywords:** epigenetic inheritance, evolution, C. elegans, small RNAs

## Abstract

Epigenetic inheritance is generally less stable across generations than DNA sequence-based heredity. One common form of epigenetic inheritance, found in plants, fungi, and animals, is the transgenerational memory of gene silencing mediated by small RNAs. These small RNAs can be amplified through RNA-dependent RNA polymerases, thus maintaining the silenced state across multiple generations. Such molecular mechanisms raise questions regarding their natural variation and evolutionary impact. We here ask whether the presence and duration of epigenetic inheritance display genetic variation within a species. We use the ability of the nematode Caenorhabditis elegans to silence genes for multiple generations after an initial RNA interference trigger. We find that the presence and the duration of silencing in number of generations differ across C. elegans wild strains. Strikingly, several wild strains show no memory, while some display a longer memory than the reference strain. Natural DNA sequence polymorphisms, such as in the set-24 and drh-1 genes, affect epigenetic memory of an external trigger, demonstrating that intraspecific DNA sequence evolution affects the duration of epigenetic inheritance. We further show that the duration of silencing memory in wild strains is quite robust to environmental variation such as diet and passage through larval diapauses, but not to temperature variation. Altogether, these results demonstrate intraspecific diversity in the regulation of small RNA-based heredity, a prerequisite for selection acting on genetic variants affecting epigenetic inheritance duration.

## Introduction

Various molecular mechanisms of epigenetic inheritance have been uncovered over the last decade across biological kingdoms, thus renewing the interest about the importance of such non-conventional inheritance in evolution (Lind and Spagopoulou, 2018; Bonduriansky and Day, 2018; Adrian-Kalchhauser et al., 2020; Duempelmann, Skribbe et al., 2020; Ashe et al., 2021). Two features distinguish epigenetic variation from DNA sequence variation: 1) epigenetic variants may originate not only stochastically but also through environmental induction; 2) epigenetic variants are generally less stable over generations than DNA sequence. We here focus on this latter point, regarding the stability of epigenetic variants across generations. At a large macroevolutionary scale, this duration ranges from long-term stability of methylated epialleles in plants (Cubas et al., 1999; Weigel and Colot, 2012; Miska and Ferguson-Smith, 2016; Fitz-James and Cavalli, 2022) or of paramutation in Drosophila (de Vanssay et al., 2012) to a lack of clear demonstration of a multigenerational memory in mammals beyond parental or grandparental effects (Heard and Martienssen, 2014). In the midst of this stability spectrum lies small RNA silencing in the nematode Caenorhabditis elegans, with a few generations of memory of the initial trigger in a commonly used experimental paradigm. This intermediate range of a few generations offers a particularly favorable case to study its variation within the species and the possible evolutionary significance of this variation.

Specifically, the C. elegans reference strain N2 is able to transmit the memory of silencing by double-stranded RNA, for example after an initial trigger of exogenous RNA interference (RNAi). The primary small RNAs are amplified in secondary small RNAs by RNA-dependent RNA polymerases (Sijen, Fleenor, Simmer et al., 2001; and Fire, 2007; Vasale, W. Gu et al., 2010; Pak et al., 2012; Tsai et al., 2015; X. Chen, K. Wang, Mufti et al., 2024). This amplification in principle can maintain the memory of the initial signal over several generations even after it has been removed (Gu et al., 2012; Billi et al., 2014; F. Xu, X. Feng et al., 2018; Shukla et al., 2021; Ouyang et al., 2022). Many factors required for the inheritance of small RNA pools were discovered using laboratory genetic screens in the N2 reference background (e.g. Grishok et al., 2000; Vastenhouw et al., 2006; Buckley, Burkhart et al., 2012; Lev, Seroussi et al. 2017; Spracklin et al., 2017; Du, Shi et al., 2023; Chen and Phillips, 2024). The RNAi memory mechanism is distinct from the RNAi mechanism itself (Grishok et al., 2000; Buckley, Burkhart et al., 2012).

This molecular mechanism of amplification and inheritance along generations raises questions regarding its ecological and evolutionary context. A prerequisite for studying how natural selection may act on epigenetic inheritance duration is to detect intraspecific diversity in the initiation and duration of epigenetic inheritance. The aim of the present work is to measure intraspecies variation for RNAi memory duration in C. elegans. Note, that we will not explore the natural variation in epialleles (here small RNA sequence representation), nor which external ecological factors may trigger initial variation in small RNA pools (Baugh and Day, 2020), but how long the memory of such triggers remains.

We probe natural strains of C. elegans for their transmission of an initial RNAi trigger and find that they differ in both occurrence and duration of their RNAi memory. We show that natural DNA sequence polymorphisms in the set-24, drh-1 and eri-6/7 genes may affect epigenetic memory of silencing. In addition to genetic variation, we find that the RNAi memory duration varies with temperature and is quite insensitive to our other tested environments, including starvation, passage through dauer diapause or bacterial diet. More importantly, the detection of intraspecific genetic variation for the presence and duration of RNAi memory means that, like any quantitative trait, epigenetic inheritance can respond to standard natural selection.

## Results

### The duration of RNAi silencing memory varies among C. elegans wild isolates

We first selected genetically divergent C. elegans wild isolates that are sensitive to RNAi. Based on the data by (Paaby et al., 2015; Crombie et al., 2019) and our prior work with the MY10 strain (Frézal et al., 2018), we focused on the eight wild strains shown in Fig. 1A. We used two distinct transgenes to assay small RNA silencing and two methods to introduce them into wild genetic backgrounds. First, we introduced a single-copy pie-1p::GFP::H2B::pie-1 transgene by repeated backcrosses to the target wild isolate background (Ashe, Sapetschnig et al., 2012). With this transgene, fluorescence is restricted to the germline nuclei and strictly nuclear because of the histone protein fusion (Fig. 1B). Second, we used CRISPR/Cas9 genome editing to introduce at the same site in all wild isolate backgrounds a specifically engineered mex-5p::ce-GFP::tbb-2 transgene, optimized for high expression and devoid of piRNA sites (Aljohani et al., 2020). As a result, the germline-specific localization of this C. elegans enhanced GFP (ce-gfp) is brighter, and fluorescence also present in the cytoplasm because, despite the nuclear localization signals, the protein is less retained in the nucleus than with the histone fusion (Fig. 1B).

**Figure 1:**
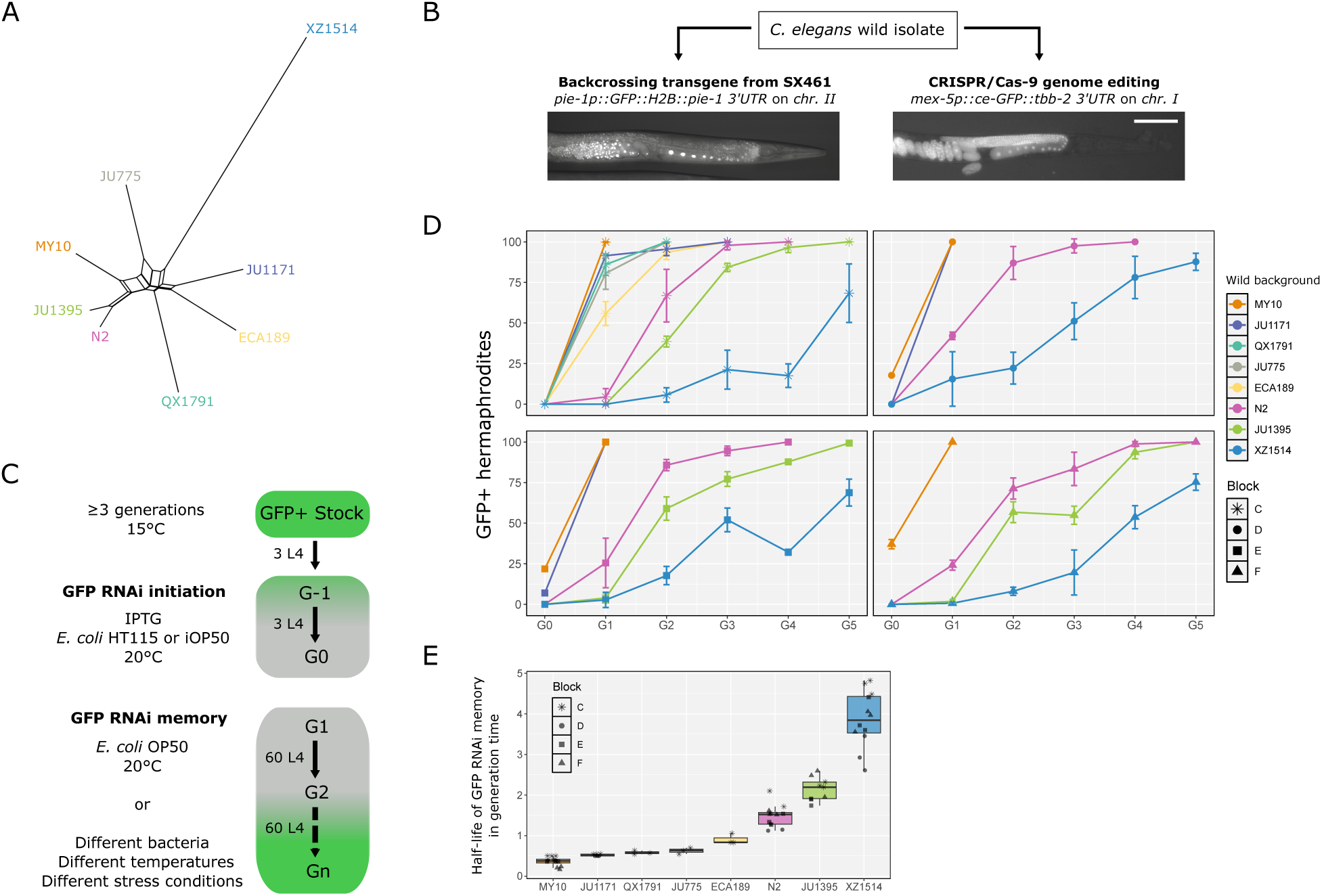
**Variation in the duration of RNAi memory among C. elegans natural isolates**. (**A**) Haplotype network of the C. elegans natural isolates used in this work. The network is based on genome-wide single-nucleotide polymorphisms from CaeNDR (Crombie et al., 2019) and built using SplitsTree. (**B**) Introduction of germline-expressed transgenes into C. elegans natural isolates, either through introgression by successive backcrosses or introduction at the same genomic site by CRISPR/Cas9-mediated editing. The two transgenes differ in cis-regulatory sequences, coding sequences and 3’UTR. The pie-1p transgene drives expression of GFP fused to histone H2B. The mex-5p transgene drives expression of a ce-GFP sequence with two NLS sequences and is devoid of predicted piRNA sites. The fluorescence micrographs show their different patterns of GFP expression. Bar: 100 µm for both pictures. (**C**) Schematic of the GFP silencing memory assay. RNAi is triggered by feeding a stock of animals expressing GFP (as indicated by the green color) with E. coli bacteria expressing double-stranded RNA against the corresponding GFP sequence for two generations (called G-1 and G0). The two transgenes strongly differ in coding sequences, and thus distinct RNAi clones were used to silence each transgene. Experimental variations during RNAi initiation or the memory assay are indicated. (**D**) Four independent experiments assaying RNAi memory variation in different isolates with the mex-5p::ce-GFP::tbb-2 transgene inserted on chromosome I. We call each of these four independent experiments a block, lettered here C, D, E and F. Three replicates were run for each line for each block (see Table S2 for raw results). In these experiments, RNAi was initiated with E. coli HT115 bacteria transformed with pMNK25. For the sake of simplicity, we indicate the name of the wild isolate background rather than that of the genome-edited strain (see Table S1 for the corresponding strain name). The rank order of the strains is highly reproducible. On the graph, the lines follow the means of the three replicates and error bars represent their standard deviation (SD). (**E**) Boxplot showing the estimated half-lives of gfp silencing memories extracted from the data in (D) (see Table S3 for raw values). Generalized linear mixed-model (glmm) statistics based on these values (excluding strains tested once) demonstrated a strong effect of the strain on the dynamics of gfp desilencing over generations: p < 2x10^-16^.

Using the protocol from (Ashe, Sapetschnig et al., 2012; Lev, Seroussi et al., 2017) (Fig. 1C), we assayed the duration of RNAi memory in the reference N2 background in parallel with the pie-1p::GFP and mex-5p::ce-GFP transgenes. After RNAi initiation, both transgenes were initially silenced in all animals (Fig. S1A). Once the RNAi trigger was removed, full silencing was maintained in the progeny (generation G1). Starting at the third generation, a fraction of the population re-expressed GFP and this percentage increased through generations. The two transgenes showed similar dynamics; however as can be expected from the lack of piRNA sites, the RNAi memory was maintained for a shorter time with the mex-5p::ce-GFP transgene (Fig. S1C). We further mainly used the mex-5p::ce-GFP transgene but report below on the convergent results using both methods.

Our first aim was to test whether wild genetic backgrounds differed in their RNAi memory duration. After a two-generation exposure to gfp RNAi from E. coli HT115 bacteria, we found that the mex-5p::ce-GFP transgene was fully silenced in all isolates, with the exception of some individuals of the MY10 strain, and that the duration of RNAi memory greatly varied among wild genetic backgrounds (Fig. 1D-E, Table S2, Table S3). Once the RNAi trigger was removed, a fraction of the population in the N2 reference re-expressed GFP at the first generation and this percentage increased until fluorescence was fully recovered in all individuals at the third or fourth generation. Some wild isolates, such as MY10, JU1171 or QX1791, did not, or hardly, transmit parental GFP silencing to the first generation. Other isolates, such as JU1395 or XZ1514, displayed an increased duration of RNAi memory compared to N2. The results were consistent across three replicates per experiment and four distinct experiments, which we call “blocks”, lettered C, D, E and F (Fig. 1D, Table S2). We plotted in Fig. 1E estimated half-lives of GFP silencing memory. Using this estimate, we found significant differences for GFP desilencing dynamics across wild genetic backgrounds (generalized linear mixed model or glmm, p < 2x10^-16^).

We further asked whether the variation among isolates could be reproduced using the pie-1p::GFP transgene. As expected, given the possible silencing by piRNAs of the pie-1p::GFP::H2B::pie-1 transgene, all displayed increased RNAi memory with this transgene compared to its mex-5p::ce-GFP::tbb-2 counterpart (Fig. S1D-E). Most importantly, the relative rank of RNAi memory duration of the tested wild isolates was well conserved (Fig. S1D), showing that they differ in the duration of their RNAi silencing memory.

### The set-24 DNA sequence polymorphism affects RNAi memory

We then aimed to determine whether DNA sequence polymorphisms could explain variation in RNAi memory, thus ruling out other types of hereditary variation that may influence this phenotype. For this, we leveraged our prior knowledge of natural variation in another multigenerational phenotype. The mortal germline phenotype (Mrt) is a multigenerational sterility process whereby lines become sterile after several generations (Ahmed and Hodgkin, 2000). Several mutations affecting small RNA pathways, chromatin remodeling and RNAi memory cause a temperature-sensitive sterility phenotype in the N2 background (Ahmed and Hodgkin, 2000; Buckley, Burkhart et al., 2012; Lev, Seroussi et al., 2017; Spracklin et al., 2017). Interestingly, some wild isolates cultured in laboratory conditions also display multigenerational sterility, which is counteracted by association with natural bacteria (Frézal et al., 2018; Frézal, Saglio et al. 2023). An example is the MY10 strain, which becomes sterile after only two to three generations at 25°C. (Frézal et al., 2018) showed that a partial deletion of the set-24 gene in MY10 (Fig. 2C) was a major effect variant causing a Mrt phenotype. set-24 encodes a SET-domain protein likely binding modified histones. Given the relationship between RNAi memory and the Mrt phenotype, and the lack of RNAi memory of the MY10 background (Fig. 1D-E), we asked whether this set-24 polymorphism (Fig. 2C) affected RNAi memory.

**Figure 2:**
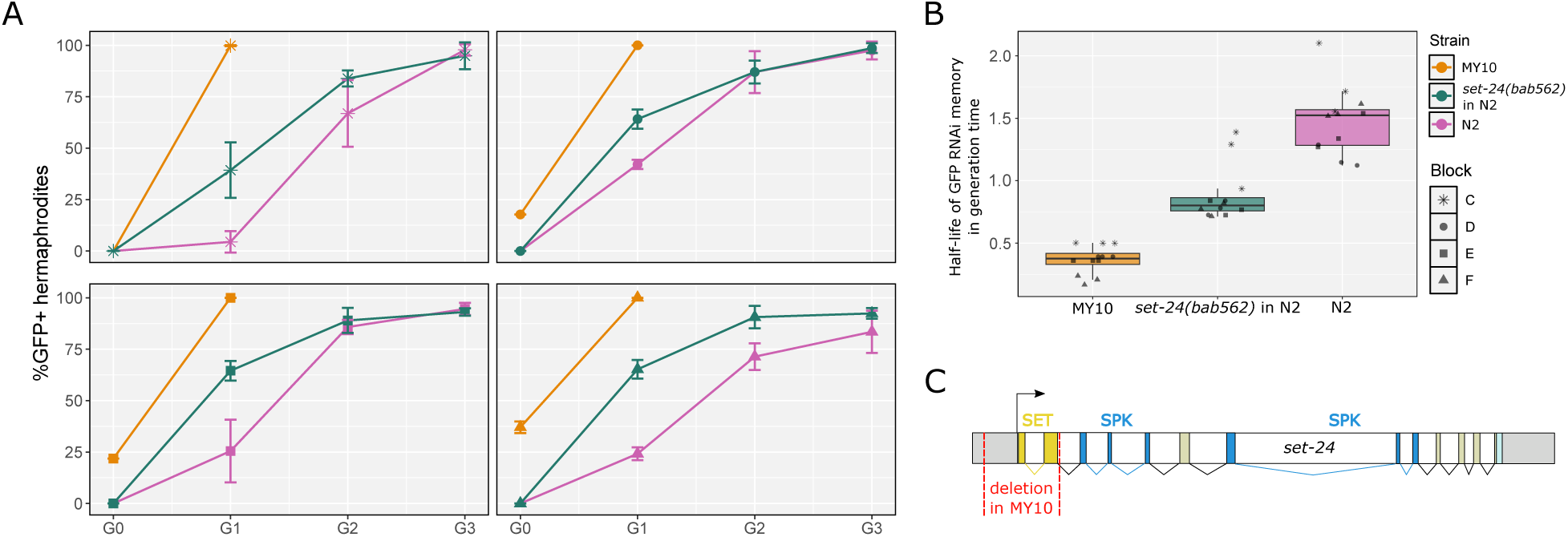
**A natural DNA sequence polymorphism in the set-24 gene affects RNAi memory**. (**A**) Effect on RNAi memory of the set-24(bab562) allele using the mex-5::ce-GFP transgene in the N2 genetic background. The four panels correspond to the same four blocks as in Fig. 2A. Lines of each graph follow the mean of three replicates and the bars represent the standard deviation of the replicates. (**B**) Boxplot showing the estimated half-lives of GFP RNAi memory. Glmm analysis based on these values followed by a Tukey pairwise comparison indicates a statistical difference for the median RNAi silencing memory between N2 and the strain carrying the set-24(bab562) allele in N2 background: p < 1x10^-4^, as well as between the latter and the MY10 strain: p < 1x10^-4^ . (**C**) Schematic depicting the set-24 gene and the location of the natural deletion in the MY10 genetic background.

For this, we engineered in the N2 background the set-24(bab562) allele that mimics the natural deletion from the MY10 isolate (called set-24(mfP23)) and crossed it to the mex-5p::ce-GFP::tbb-2 transgene. We found that the set-24(bab562) allele significantly reduced RNAi memory compared to the N2 genetic background (Fig. 2A-B). This set-24 polymorphism explained only part of the difference in RNAi memory between the N2 and MY10 backgrounds, suggesting that further polymorphisms may underlie some RNAi memory difference between the two strains. Altogether, this result demonstrates that DNA sequence evolution affects epigenetic memory.

### The intermediate frequency drh-1 polymorphism affects RNAi memory

Given that the derived set-24 deletion allele is rare in the wild (Frézal et al., 2018), we wondered whether more common and thus potentially more evolutionary stable polymorphisms could underlie natural variation in the duration of RNAi memory. C. elegans antiviral defense mechanisms rely in part on recognition of viral double-stranded RNA (dsRNA) and production of small RNAs targeting the virus (Ashe, Bélicard, Le Pen, Sarkies et al., 2013; Coffman et al., 2017; Sowa et al., 2020; Batachari et al., 2024). As the machinery required to recognize and initiate antiviral response is in part the same as that of RNAi (Tabara et al., 2002; Consalvo et al., 2024), we wondered whether polymorphisms found in the double-stranded RNA antiviral pathway could also affect RNAi memory.

One gene involved in the complex responsible for foreign dsRNA recognition and degradation is drh-1, a RIG-I like DExD/H box helicase with dsRNA binding activity (Duchaine et al., 2006; Batachari et al., 2024; Consalvo et al., 2024). Studying variation in the sensitivity to Orsay virus infection among wild C. elegans strains, we previously found that a 159 bp natural deletion allele (niDf250) in the RIG-I like domain of DRH-1 was causally associated with hypersensitivity to viral infection and was present at an intermediate frequency in the species (Ashe, Bélicard, Le Pen, Sarkies et al., 2013) (Fig. 3A).

**Figure 3:**
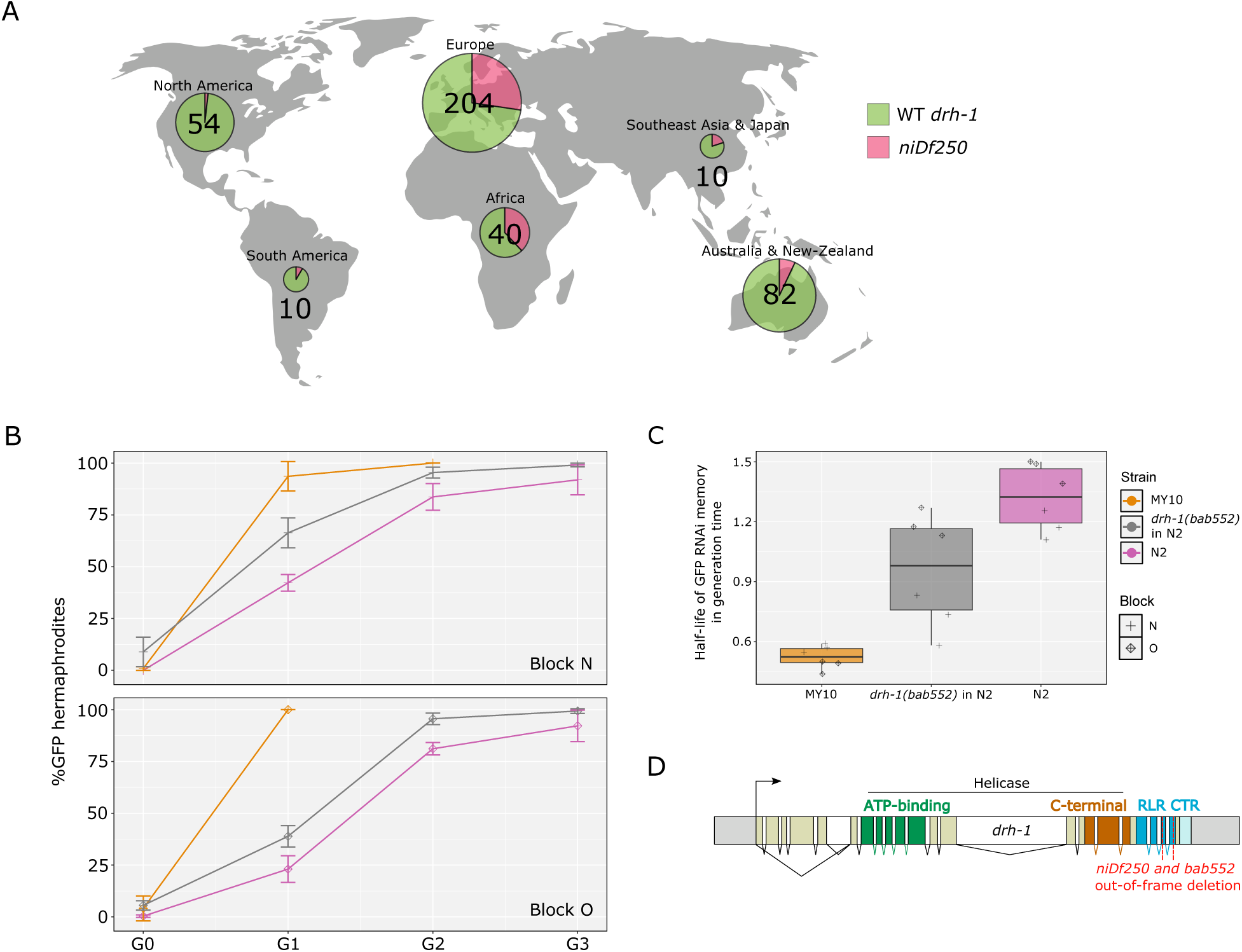
**A common natural DNA sequence polymorphism in the drh-1 gene affects RNAi sensitivity and memory**. (**A**) Diagram depicting the proportion of isotypes possessing the 159 bp deletion in drh-1 by world region. Numbers shown correspond to the number of isotypes sequenced by continent from CaeNDR. As no isolate sampled in Hawaii presents this polymorphism, Hawaii was excluded from the representation. Created with BioRender.com. (**B**) Effect on RNAi memory of the drh-1(bab552) allele using the mex-5::ce-GFP transgene in the N2 genetic background. Panels correspond to blocks N and O. In these two blocks, E. coli iOP50 bacteria transformed with pFF1 (see Methods) were used to initiate RNAi against gfp. Lines of each graph follow the mean of three replicates and the bars represent the standard deviation of the replicates. (**C**) Boxplot showing the estimated half-lives of GFP RNAi memory of the strains tested in blocks N and O. Glmm analysis based on these values followed by a Tukey pairwise comparison indicates a statistical difference for the median RNAi silencing memory between N2 and drh-1(bab552) in N2: p = 1.1x10^-3^; between MY10 and drh-1(bab552) in N2: p < 1x10^-4^.

To test whether this drh-1 allele affected RNAi memory, we reproduced this natural deletion by genome editing of the N2 genetic background, yielding allele drh-1(bab552), and crossed it to the mex-5p::ce-GFP::tbb-2 transgene. The RNAi efficacy was tested in this line along with the N2 and MY10 backgrounds, the latter a natural carrier of the deletion. We first observed that initiating RNAi using E. coli iOP50 transformed with pFF1 (see Methods) increased RNAi efficiency as even MY10 was almost fully silenced at G0 (Fig. 3B, Fig. S4). We also noted that, when placed in an N2 background, the partial deletion of drh-1 not only reduced gfp RNAi memory but made animals somewhat less responsive to RNAi initiation (Fig. 3B-C). The drh-1 deletion allele is maintained at intermediate frequency in the C. elegans species, particularly in Eurasia and Africa and is thus a common allele potentially affecting RNAi memory (and marginally, efficiency) in natural populations.

### Structural polymorphisms at the eri-6/7 locus likely affect RNAi memory

Besides exogenous double-stranded RNAs, different endogenous small RNA pathways exist in C. elegans. In both cases, the primary small RNAs are amplified in secondary 22G small RNAs (Billi et al., 2014). Because factors required for the amplification of exogenous and endogenous small RNAs are shared, the different small RNA pools compete amongst each other for amplification (Houri-Ze’evi et al., 2016; Shukla et al., 2021; Karin et al., 2023). For instance, the endogenous piRNA pathway modulates the duration of exogenous RNAi memory: the inactivation of the piRNA pathway results in a longer exogenous RNAi memory (Shukla et al., 2021). Inactivating mutations in the ERGO-1 26G-RNA pathway enhance RNAi (Eri phenotype) (Kennedy et al., 2004; Lee et al., 2006; Fischer, Butler et al., 2008) but a potential effect on RNAi memory had not been tested.

The eri-6/7 locus produces a helicase involved in the endogenous ERGO-1 26G-RNA pathway repressing retrotransposons and novel genes (Fischer et al., 2011; Fischer and Ruvkun, 2020). The gene has been shown to greatly vary in structure among C. elegans wild isolates (Fig. 4C), with an inversion of the eri-6 part of the gene and further indels, impacting the splicing efficiency of the inverted eri-6 exons to the remaining eri-7 exons and the expression of their downstream targets (Fischer, Butler et al., 2008; G. Zhang et al., 2024).

**Figure 4:**
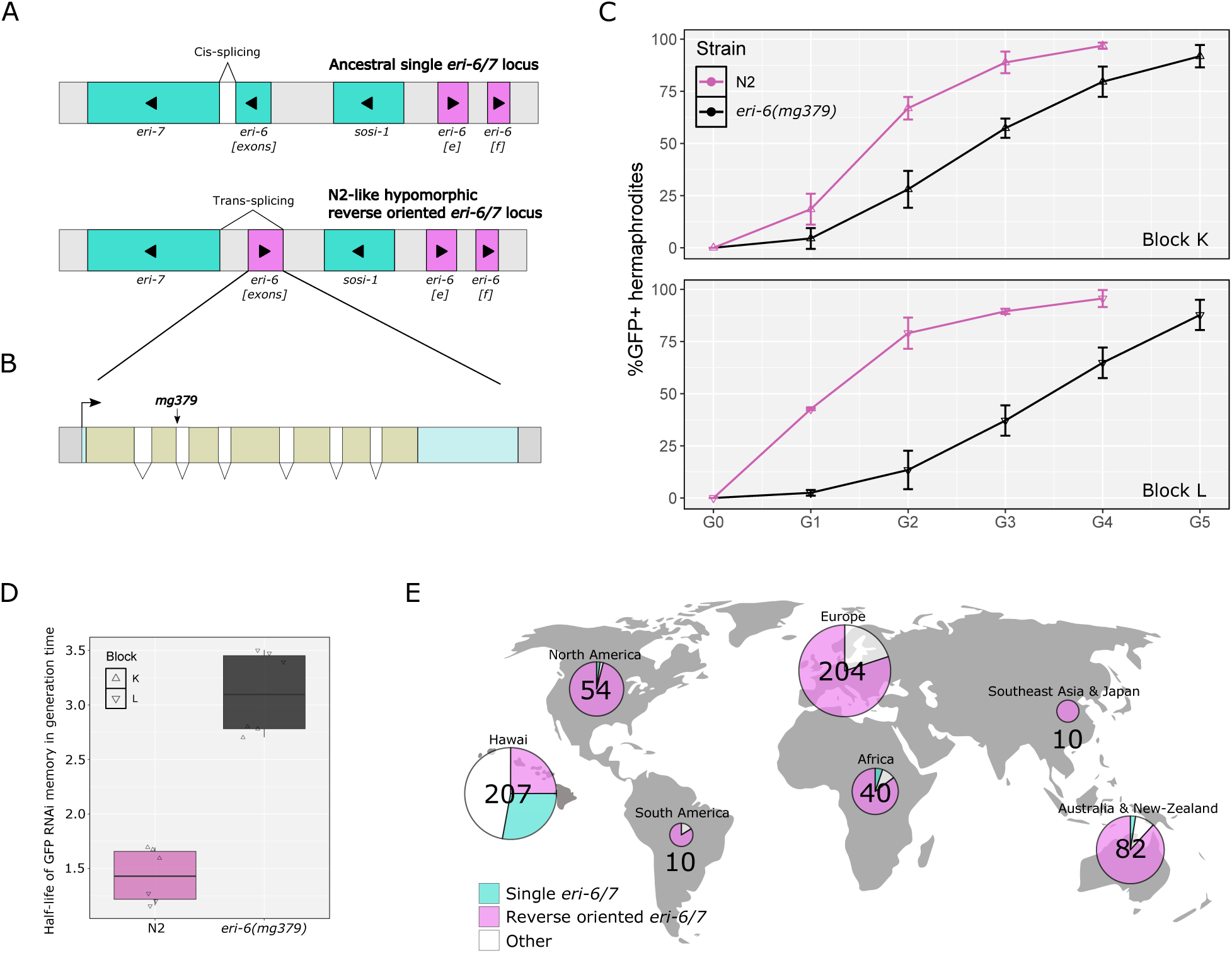
**Competition between endogenous and exogenous small RNA pathways alters RNAi memory duration**. (**A**) Diagrams depicting two commonly found gene structures of the eri-6/7 locus in C. elegans wild strains (here additional introns within each large block are omitted). The ancestral state of this locus harbors a unique gene in the right to left orientation (colored in blue). The more recent N2-like structure of eri-6/7 is thought to have arisen from a transposable element invasion, a resulting inversion of the first exons (pink orientation, left to right) forming the so-called eri-6 part of the gene. This rearrangement of eri-6 is predominant in isolates outside of Hawaii and is hypomorphic (Fischer, Butler et al., 2008; G. Zhang et al., 2024). (**B**) Diagram showing the detailed gene structure of eri-6 as found in the N2 genetic background. The laboratory mg379 allele is a splice-donor mutation. (**C**) RNAi memory assay (blocks K and L) of the eri-6(mg379) mutant, using the mex-5p::ce-GFP transgene. This eri-6 reduction-of-function mutation decreases endogenous small RNA production, thus enabling a better amplification of 2° siRNAs corresponding to exogenous dsRNAs and leads to a longer RNAi memory. E. coli iOP50 was used to initiate gfp RNAi. (**D**) Boxplot showing estimated half-lives of GFP RNAi memory for N2 and eri-6(mg379) in blocks K and L. Glmm on these half-life values followed by a Tukey pairwise test indicates a statistical difference of median RNAi silencing memory between strains: p < 1x10^-4^. (**E**) World map presenting the percentage of isotypes possessing either one of the eri-6/7 gene structures shown in (A) relative to the total number of isotypes by world region in CaeNDR. Other structures of this locus comprise, for example, a single eri-6/7 gene without sosi-1 and eri-6[e][f] genes, eri-6[exons] surrounded by remnants of the transposable element and an N2-like structure with duplicated eri-6[exons (G. Zhang et al., 2024). Created with BioRender.com.

We tested whether the laboratory-induced eri-6(mg379) splice site mutation (Fig. 4B) affected the duration of exogenous RNAi silencing memory. We found that this reduction-of-function mutation increased RNAi memory (Fig. 4A), possibly through the competition between small RNA pools for amplification. The natural rearrangements impacting the eri-6/7 locus activity thus likely affect RNAi memory in C. elegans.

### Environmental effects on RNAi memory of C. elegans wild isolates

Having demonstrated a genetic component in the duration of GFP silencing, we investigated whether the environment may impact this duration. In natural contexts, C. elegans follows a boom-and-bust life cycle (Fig. 5A) and is subject to multiple biotic and abiotic constraints (Félix and Duveau, 2012; Frézal and Félix, 2015; Schulenburg and Félix, 2017). We focused on some of these relevant factors to assess their potential impact on RNAi memory: temperature, passage through diapause states (Fig. 5B-G), and bacterial environment (Fig. S2A). In order to keep RNAi efficiency constant across environmental conditions, we initiated GFP silencing memory as above until G0 and modified the conditions in which the subsequent generations were cultured.

**Figure 5:**
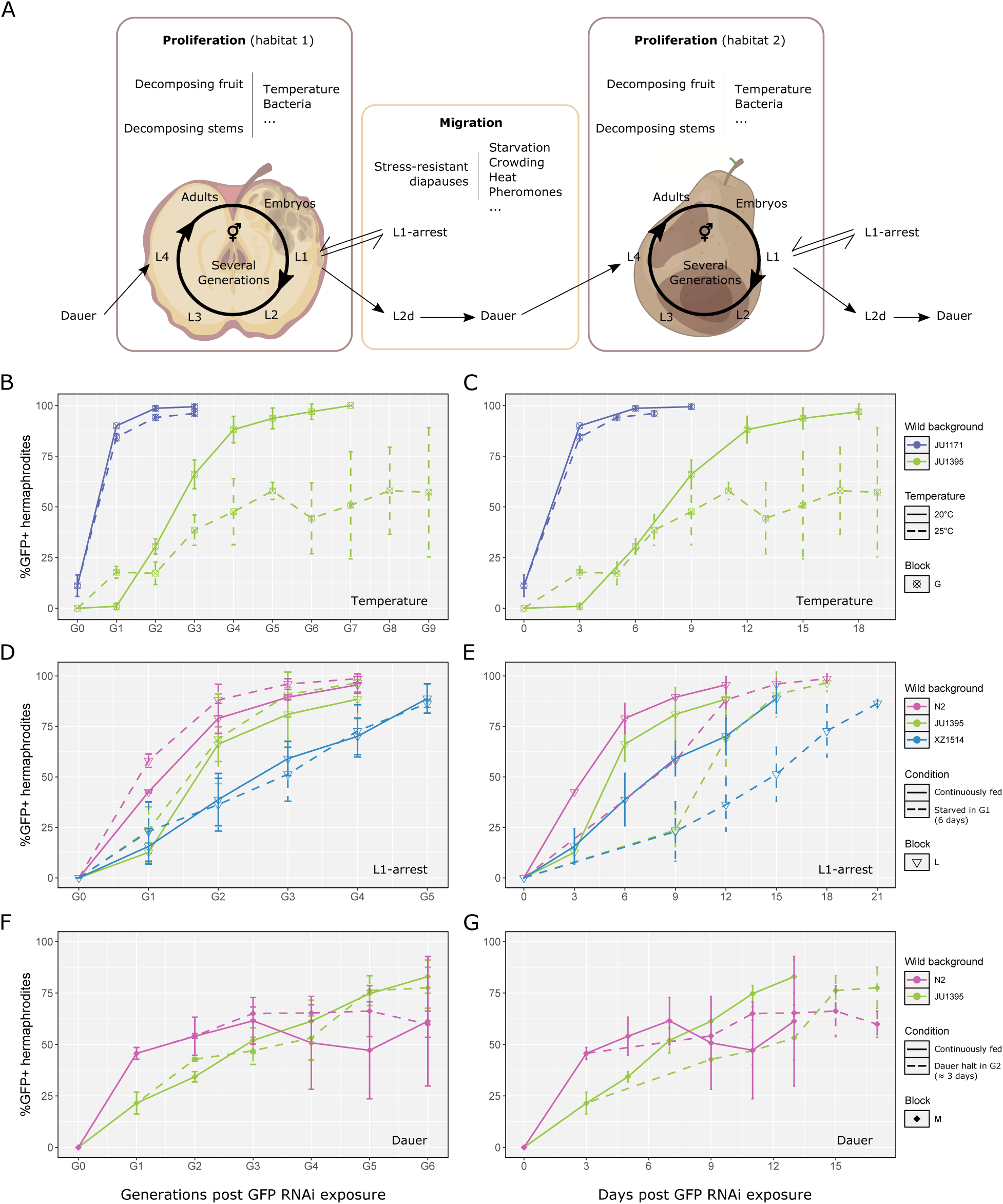
**Impact of the environment on the duration of RNAi memory**. (**A**) Schematic illustration of the “boom-and-bust” life cycle of C. elegans natural populations (created with BioRender.com). (**B-G**) Tests of temperature [B-C], L1 arrest [D-E] and dauer diapause [F-G] on RNAi memory duration using the mex-5p::GFP transgene, initiating RNAi against gfp with E. coli iOP50 bacteria transformed with pFF1. The results are plotted along the number of generations in [B, D, F] and number of days in [C, E, G]. For each block and condition, 3 replicates were run per strain. Graph lines represent means and bars the standard deviation. (**B-C**) Effect of temperature on RNAi memory in the JU1171 and JU1395 backgrounds (block G). Culturing JU1395 at 25°C increases duration of its RNAi memory in number of generations (glmm on half-lives of GFP RNAi memory in blocks C and G, p < 1x10^-4^) and in days (p = 5.3x10^-3^). No significant difference was observed for JU1171. See Fig. S2B for an additional experiment using the N2 and JU1395 backgrounds. (**D-E**) Effect of L1 arrest for 6 days at 20°C at the first generation after parental RNAi exposure for strains in the N2, JU1395 and XZ1514 backgrounds (block L). Analyzing the half-lives of the GFP RNAi memory of N2 and JU1395 in blocks K and L in a glmm framework, a significant reduction between control and starvation conditions were found in generations, p = 2,8x10^-3^ and 5x10^-4^, for N2 and JU1395, respectively, as well as a significant extension of RNAi memory in absolute time, p < 1x10^-4^ for each strain. (**F-G**) Effect of a ∼3-day long dauer diapause at the second generation after GFP RNAi exposure (block M). No effect was found on GFP de-silencing dynamics when expressing the results in number of generations (glmm on half-lives, followed by Tukey’s test, p > 0.45 between fed and dauer condition for both N2 and JU1395). However, the RNAi memory was extended in number of days after passage through dauer in G2 in the JU1395 background (p = 5.6x10^-3^ between fed and dauer conditions). This experiment was performed at 25°C.

We first compared RNAi memory at two temperatures, 20°C (temperature of previous assays) and 25°C, on two strains displaying different memory durations. We found that the JU1395 background strain transmitted GFP silencing for a higher number of generations at 25°C than at 20°C, whereas the higher temperature did not rescue the lack of memory of the JU1171 strain (Fig. 5B-C). Development occurs faster at 25°C compared to 20°C, therefore we tested whether the longer memory in number of generations corresponded to a similar memory in absolute time. Instead, the JU1395 background still displayed a longer memory at high temperature in absolute time (Fig. 5B-C). We further tested the RNAi memory of the N2 strain at 25°C versus 20°C. Similar to the JU1395 background, culturing N2 at the higher temperature resulted in an enhanced gfp RNAi memory, in number of generations and in absolute time (Fig. S2B).

When facing harsh conditions during post-embryonic development, C. elegans can arrest developmentally, and survive for several weeks without eating or reproducing (Hu, 2007; Baugh, 2013); when encountering new favorable conditions, normal growth can be restored (Fig. 5A). For example, the L1 larval arrest arises after hatching in the absence of food (Baugh, 2013). To reproduce this L1 developmental arrest, we let the first generation after GFP RNAi exposure hatch in NGM plates without E. coli and remain for six days at 20°C, as in Houri-Zeevi et al. (2021). L1 larvae were then fed again in standard conditions and RNAi memory assayed. This early L1 arrest slightly reduces RNAi memory for the three tested strains when counting in number of generations (Fig. 5D). Due to the time spent in L1 arrest, when plotting in absolute time, RNAi memory was increased (Fig. 5E).

The stress-resistant dauer diapause, commonly found in C. elegans natural populations (Schulenburg and Félix, 2017), corresponds to a specialized form of the early third larval stage (Hu, 2007). Experimentally, dauer larvae can be obtained in conditions of crowding, paucity of food and elevated temperature on young larvae (Karp, 2018). In order to keep the conditions of RNAi initiation constant, we induced dauer larvae by letting the first generation (G1) animals lay embryos without transfer to a new food source and selected for dauers after a few days (see Methods). Passage through dauer for 5 days in the second generation after RNAi initiation did not alter RNAi memory kinetics of the N2 and JU1395 strains (Fig. 5F). Again, taking into account the diapause delay, passage through dauer increased RNAi memory in absolute time (Fig. 5G). We further tested the long memory background XZ1514 with dauer induction at 20°C, reproducing this procedure at every generation after G2, and similarly did not see an effect of passage through dauer (Fig. S2C). Thus, we conclude that the dauer stage, like the early L1 arrest, does not strongly affect memory in number of generations and is able to maintain the memory of RNAi through the days of diapause.

Bacteria that are found naturally associated with C. elegans encompass many different genera (Dirksen et al., 2016; Samuel et al., 2016). We tested four such bacterial wild isolates, Leucobacter CBX151, Chryseobacterium JUb044, Acinetobacter BIGb0102 and Comamonas BIGb0172, selected for their ability to sustain C. elegans growth for multiple generations while being potential pathogens (Hodgkin et al., 2013; González and Félix, 2024). We found that RNAi memory of the JU1395 and JU1171 wild backgrounds was conserved relative to E. coli OP50, when cultured with these naturally-associated bacteria. An exception is the JU1395 strain, which when tested on Leucobacter CBX151 displayed a slight reduction in memory duration (Fig. S2A).

## Discussion

We here provided evidence for microevolutionary variation in small RNA inheritance in C. elegans. The duration of epigenetic inheritance can be treated as a quantitative trait in theoretical (Helanterä and Uller, 2010; Day and Bonduriansky, 2011) and experimental models of selection.

### Natural genetic variation in RNAi inheritance

Like RNAi efficiency itself (Pollard and Rockman, 2013; Chou et al., 2024), the small RNA inheritance trait is likely to have a polygenic basis. We provide evidence for polymorphisms in several genes being involved, potentially acting at distinct points in the inheritance mechanism. These may correspond to different parameters in mechanistic models of small RNA memory, such as that developed by (Karin et al., 2023), which models feedback amplification between small RNA and histone modifications, a resulting variation in mRNA level, and a queuing system of small RNAs. Different molecular mechanisms for the extinction of memory may be at stake as well (Shukla et al., 2021; Knudsen-Palmer et al., 2024). Furthermore, after RNAi initiation, two phases have been experimentally distinguished in transgenerational silencing, based on the mutation effect of different genes in the N2 background (Woodhouse et al., 2018): establishment of RNAi inheritance at the first generation and maintenance in further generations (Fig. S3).

The drh-1 deletion allele seems to mildly diminish sensitivity to germline RNAi and severely prevents RNAi inheritance (Fig. 3B, Fig. S4A). Memory may be simply affected by the slightly lesser efficiency of the initial trigger, resulting in a less efficient initial amplification. Alternatively, competition between small RNA pools along the generations may be at stake, as seen with piRNAs or endosiRNAs, or as modeled by (Karin et al., 2023). DRH-1 is a cytoplasmic protein (Batachari et al., 2024), expressed in both somatic and germline cells (Wu et al., 2012). It recognizes preferentially blunt dsRNA (Consalvo et al., 2024) and is involved in viral RNA recognition as well as a small RNA response to mitochondrial stress (Mao et al., 2020). In an evolutionary context, drh-1 polymorphism may be driven by these different phenotypes alongside its effect on RNAi memory.

The set-24 gene codes for a SET and SPK-domain protein, recently characterized by Zeng, Furlan, Almeida et al. (2025). The SET-24 protein is localized to the germline nuclei. Through its interaction with the epigenetic regulator HCF-1 and chromatin remodeling complexes, SET-24 regulates H3K4me3 levels of hundreds of genes. Its dysregulation perturbates the pools of inherited small RNAs, which leads to impaired inheritance and a shorter RNAi memory (Zeng, Furlan, Almeida et al., 2025).

The eri-6/7 locus quantitatively affects the maintenance of the memory, likely by competition between pools of small RNAs, as modeled by Karin et al. (2023), whereby amplification of endo-siRNAs requiring ERI-6/7 competes with other small RNA pools. The partial inversion of the eri-6/7 locus n the N2 background results in a hypomorphic allele (G. Zhang et al., 2024), thus likely increasing RNAi memory. We note however that the only strain with an ancestral single orientation eri-7 locus in our set, XZ1514 (G. Zhang et al., 2024), has a long memory so other variants elsewhere in the genome of this strain likely affect the trait. It is likely that the variation in duration of small RNA memory is highly polygenic in C. elegans wild strains.

### Environmental effects on RNAi inheritance

Given the boom-and-bust lifecycle of C. elegans with a dispersal between food patches in which the environment may differ (Félix and Duveau, 2012), we expected that starvation and passage through the dispersal dauer stage may erase the small RNA memory. However, of the different culture environments we tested during the memory phase, including dauer, only temperature variation resulted in a large change, with an enhanced RNAi memory at 25°C compared to 20°C. This effect of temperature appeared after one generation of initiation of the memory.

Houri-Zeevi et al. (2021) instead suggested that small RNA memory was ‘reset’ by different stressing environments. They found a shortened memory, from the first generation, after a transient heat shock (37°C). In contrast, they had previously shown that inheritance of silencing relied on the activity of the heat-shock transcription factor HSF-1 (Houri-Zeevi et al., 2020), an effect that could be relevant in our experiments. hsf-1 overexpression leads to suppression of endogenous small RNA (sRNA) pathway components and, likely due to competition among sRNA pools, a higher proportion of silenced progeny. Using a 25°C temperature as a trigger rather than an environment during the memory phase, others studies showed that this temperature altered siRNA pools for 3-4 generations (Schott et al., 2014; Belicard et al., 2018) and de-repressed a silenced multi-copy fluorescent reporter for over 10 generations (Klosin et al., 2017). Temperature is thus an ecologically relevant factor initiating and modulating small RNA memory.

Concerning L1 stage starvation, Houri-Zeevi et al. (2021) reported a ‘resetting’ of silencing memory, analyzing statistically each successive generation as if independent. Following the same experimental procedure, and analyzing instead half-lives of silencing, we observed a statistically significant yet mild difference on RNAi memory of C. elegans between fed and starved conditions. Moreover, after halting development in the L1 or dauer stages, silencing is maintained for a longer time in number of days. Knowing that C. elegans in the wild remains in the dauer stage possibly for weeks in between nutrient-rich sources, questions arise regarding the ecological relevance of memorizing conditions of a previous environment.

### Conclusions and perspectives

A transient epigenetic memory may be advantageous in environments that change on the same timescale, with correlation between successive environments (Jablonka et al., 1995; Lachmann and Jablonka, 1996; Rivoire and Leibler, 2014). In that case, the frequency of environmental change and the life cycle duration of the organism will determine whether multigenerational memory is advantageous. One can imagine the evolution of different memory durations, with cases relying more on plasticity within a generation than on inheritance. Using C. elegans experimental evolution, (Dey et al., 2016) showed that parental effects could be selected in sequences of environments that vary at every generation. Our findings of polymorphisms that affect small RNA inheritance will allow to test which environmental sequences may favor one or the other allele. In addition, polymorphisms affecting RNAi memory duration may be selected for a pleiotropic effect on another trait. The prevalence of different forms of non-genetic inheritance across species highlights the importance of evaluating experimentally how it itself evolves and might impact the course of evolution.

## Materials and methods

### C. elegans culture and strains

C.IZelegans was cultured on 551mm NGM agar plates (Stiernagle, 2006) seeded with 100 μl of a saturated culture of E.IZcoli OP50 grown overnight in Luria Broth (LB) at 37°C, except if otherwise indicated. C.IZelegans wild isolates were obtained from the Caenorhabditis Genetics Center (CGC), Michael Ailion, Erik Andersen, Leonid Kruglyak or our own collection. The set-24(bab562) and drh-1(bab552) mutants were engineered using CRISPR/Cas9 editing by the CNRS SEGICel facility (Lyon, France), mimicking the natural alleles characterized in Frézal et al. (2018) and Ashe, Bélicard, Le Pen, Sarkies et al. (2013), respectively. The eri-6(mg379) mutant (Fischer et al., 2008) was obtained from the CGC. Bleaching of cultures was performed as in Hibshman, Webster et al. (2021).

### E. coli strains and plasmids

The gfp targeting sequence directed against the pie-1p::GFP transgene was obtained from Eric Miska’s laboratory and inserted into the L4440 vector. The gfp targeting sequence directed against the mex-5p::ce-GFP transgene was inserted into plasmid L4440 (Timmons et al., 2001), yielding plasmid pMNK25. The NotI-KpnI digested insert was subcloned into the T444T plasmid (Sturm, Saskoi et al., 2018), yielding plasmid pFF1, the latter harboring transcriptional terminators (Table S4). The RNAi clone that was used is indicated for each experiment. In our first experiments, E. coli bacterial strains HT115 was used transformed with plasmid pMNK25 and in later experiments, E. coli iOP50 was used with pFF1. The latter combination seems to result in a stronger RNAi efficiency, but the overall variation among strains is comparable for all clones.

### Bacterial strains

The non-E. coli bacteria were originally isolated from different Caenorhabditis collected in the wild. CBX151 Leucobacter was found infecting a Caenorhabditis collected in Cape Verde (Hodgkin et al., 2013). JUb044 Chryseobacterium was isolated from Santeuil, France; BIGb102 Acinetobacter and BIGb172 Comamonas were collected in decomposing apples in Orsay, France (Samuel et al., 2016). These bacteria were grown in LB liquid culture for 1 16 or 1 32 h with 220 rpm agitation at 22°C for CBX151 or 28°C for the other bacteria.

### Introgression of the pie-1p::GFP::H2B::pie-1 transgene

We initially used the mjls31[pie-1p::GFP::H2B] transgene (Ashe, Sapetschnig et al., 2012), introduced by MosSCI (Frøkjær-Jensen et al., 2008) in the reference strain N2 on chromosome II at 8.4 Mb, yielding strain SX461 (Ashe, Sapetschnig et al., 2012). As this transgene is easily silenced, we first desilenced its expression using mut-7 RNA interference. We then introgressed the transgene in several wild isolates by at least six rounds of backcrossing to the target wild genetic background, using SX461 males in the first cross. The primers used to verify the genotype of the different chromosomes are in Table S5 for JU1171, MY10 and JU775, with specific attention to the presence of set-24(mfP23)II in the MY10 genetic background; genotyping was not performed for JU1395 because of its close relatedness to N2. As this transgene was introduced in different wild isolates through backcrosses (Ashe, Sapetschnig et al., 2012), a N2 genomic segment linked to it is also present and of a different length for each backcross. As these N2 genomic regions could in principle affect RNAi memory compared to the target wild genomic background, we also used CRISPR/Cas-9 genome editing to introduce the transgene at a given locus, circumventing this problem.

### Introduction of the mex-5p::ce-GFP::tbb-2 3’UTR transgene by genome editing

A CRISPR-Cas9 protospacer was selected (TCCGTGTCTTACTACTGTA) in a conserved region on chromosome I that is permissive for germline expression (El Mouridi et al., 2022). The protospacer was added to an sgRNA(F+E) scaffold (Chen et al., 2013) and ordered as a gene fragment (Twist Bioscience, CA, USA). The gene fragment was subsequently cloned into an empty ampicillin resistant vector using Gibson assembly (Gibson et al., 2009) to give pMDJ344. An expression vector containing a mex-5 promoter (pCFJ645) (Zeiser et al., 2011), a nuclear C. elegans-optimized GFP (pCFJ2389) (Aljohani et al., 2020), a tbb-2 3‘UTR (pCM1.36) (Merritt et al., 2008), and a hygromycin resistance cassette (pCFJ767) (Radman et al., 2013) were cloned using a multi-site Gateway reaction (Invitrogen cat. no. 12538200) to give pMDJ343. Roughly 250 bp homology arms from either end of the selected CRISPR-Cas9 cut site were added to pMDJ343 using Gibson assembly (Gibson et al., 2009) to make the final repair template pMDJ346. The resulting lines are listed in Table S1.

A similar procedure was performed exclusively on an N2 genetic background, this time using a (CCGTGGAATCAAGTTAATC) CRISPR-Cas9 protospacer in a conserved region of chromosome V. By using the same methodology as stated above, the resulting gene fragment cloned into an empty ampicillin resistant vector gave pMDJ345. The adding of 250 bp homology arms from either end of this chromosome V cut site to the mex-5p::ce-GFP::tbb-2 transgene and its hygromycin resistant cassette gave pMDJ347. This insertion was used to cross the eri-6 mutation located on chromosome I.

Injection mix contained a codon-optimized Cas9 expression vector (pCFJ2474) (Aljohani et al., 2020) at 25 ng/µl, the sgRNA vector (pMDJ344) at 5 ng/µl, the repair template (pMDJ346) at 10 ng/µl, a linearized histamine marker (pSEM238) (El Mouridi et al., 2021) at 10 ng/µl, a pan-muscular mCherry marker (pSEM235) (El Mouridi et al., 2020) at 10 ng/µl, and a DNA ladder (1 kb plus, Invitrogen) at 40 ng/µl to make up a final mix concentration of 100 ng/µl. 10-20 young adults were injected for each wild isolate as previously described (Mello et al., 1991) and placed at 20°C. 500 µl of hygromycin (4 mg/ml) was added three days post injection to select for extrachromosomal array animals (Radman et al., 2013). Once plates starved (7 days post injection), 500 µl of histamine (500 mM) were added to paralyze extrachromosomal array animals (El Mouridi et al., 2021). Single copy insertion lines are characterized by hygromycin resistance, insensitivity to histamine, lack of somatic mCherry fluorescence, and germline ce-gfp expression.

### Comparison between the transgenes

The pie-1p::GFP transgene possesses three introns and four exons. Sequence of gfp in the mex-5::ce-GFP transgene is around 1.75 kb and possesses six introns and five exons.

The mex-5p::ce-GFP transgene was custom-built for this work. It includes an engineered gfp sequence that was designed to lack piRNA homology sites and favor transgene expression (Aljohani et al., 2020). We cannot rule out that the piRNA loci differ among the C. elegans wild isolates and that for example some of the long memory strains possess piRNAs targeting this transgene. However, we note that the same ranking of wild strains regarding RNAi memory was observed with the pie-1p::GFP transgene, which has a quite different sequence, which only matches that of the mex-5p::ce-GFP transgene on 571/792 nucleotides along the mRNA. For RNAi, we used E. coli produced dsRNAs (Table S4) with a perfect match against the length of the corresponding mRNA.

### Culture plates for RNAi interference

Empty 55 mm Petri dishes were poured with autoclaved NGM medium supplemented with a 2 µm filtered IPTG solution to obtain a final concentration of 1 mM. Plates were left to dry overnight and covered at room temperature. In parallel, liquid cultures of dsRNA vector containing-bacteria were incubated for ∼16 hrs at 37°C with agitation at 220 rpm. The next day, the bacterial liquid cultures were used to seed now solidified NGM-ITPG plates with 100 µL and left to dry overnight covered at room temperature. This process was repeated for the two generations of exposure to RNAi (G-1 and G0).

### GFP silencing memory assay

Stock populations of GFP+ nematodes were maintained for at least three generations at 15°C before starting an experiment. Three replicates were conducted in parallel for each. RNAi was initiated 20°C by randomly picking three L4 stage individuals from the stock populations onto RNAi plates. Three L4 stage larvae derived from these founders (designated generation G-1) were randomly chosen for transfer onto fresh RNAi plates. In block A, 60 L4 larvae of the next (G0) generation were transferred from the RNAi memory environment to standard conditions, without RNAi trigger. For all other blocks, the G0 adults were treated with a “bleach” solution (Hibshman et al., 2021) to break down their cuticle and isolate their embryos, thus killing dsRNA expressing bacteria. Embryos were then deposited onto standard 20°C NGM plates seeded with saturated OP50 E. coli, unless indicated otherwise. These embryos were designated as generation G1. At each generation, 60 random L4 stage individuals were passed to produce the next generation and the remaining population chunked onto a fresh new NGM-OP50 plate to avoid starvation. The next day, the chunked population was scored and the transferred individuals were removed and their progeny allowed to grow.

Changes in the environment were performed during the memory phase, thus after the initiation step of parental RNAi exposure at 20°C in G-1 and G0.

Temperature: After the initiation step, subsequent generations were cultured at 25°C or 20°C on standard NGM medium seeded with 100 µL of saturated E. coli OP50 culture.

Bacteria: After the initiation step, subsequent generations were cultured at 20°C in standard NGM medium seeded with 100 µL of saturated cultures of the tested bacteria.

Starvation: After the initiation step, the bleach-treated embryos were deposited onto an empty NGM plate and left to starve for 6 days at 20°C. At this time point, L1 larvae were collected in M9 solution, centrifuged 1 min at 3000 rpm and transferred to standard culture plates to resume their development and continue the GFP silencing memory experiment as described above.

Dauer: In order to keep the conditions of RNAi initiation constant, we induced dauer larvae by letting the first-generation animals lay embryos without transfer to a new food source. Specifically, after initially transferring 60 L4 stage G1 individuals onto new plates for the fed control, the remaining population was separated in two, half being fed for scoring of the G1 generation and the other half left on the plate. The latter was then placed at either 20°C or 25°C for 7 and 5 days, respectively, in order to obtain a mix of L1/L2 arrest and dauer larvae of the next generation. 1% sodium dodecyl sulfate (SDS) was then used to filter out dauer individuals from the rest by ∼20 min incubation with mellow agitation (Karp, 2018). Surviving dauer larvae were then deposited back onto standard culture plates to resume development and continue the assay.

### Microscopy and GFP expression quantification

Animals harboring the pie-1p::GFP transgene were scored using a Zeiss AxioImager M1 microscope with a x63 objective and a Semrock GFP-1828A-000 filter. Animals harboring the mex-5p::ce-GFP transgene were scored using a Nikon AZ100 fluorescence microscope with a 5x objective and level 3 zoom and a Semrock GFP-3035D filter. Hermaphrodite adults were collected using M9 solution and pelleted waiting ∼2 minutes. A 15 µL drop of pelleted nematodes was deposited on a microscope slide containing a thin layer of 4% agar noble gel supplemented with 100 µM of NaN_3_. Around 100 individuals were scored for each replicate using no transillumination light. During the silencing recovery period, with the pie-1p::GFP transgene, nematodes were considered “OFF” when both arms were silenced or “ON” when at least one mature oocyte presented nuclear fluorescence. With the mex-5p::ce-GFP transgene, intensity and site of GFP expression varied among individuals leading us to sort nematodes according to their GFP pattern and brightness levels into “OFF”, “DIM” or “ON” categories. OFF individuals were those showing no nuclear fluorescence in their gonad, DIM individuals were those with varied patterns of expression but no nuclear fluorescence in their proximal gonad and ON individuals expressing GFP+ meiotic oocyte nuclei. Results with these three categories are shown in Table S2 and Figure Sn. The OFF and DIM categories were grouped for plotting and analysis as we considered that DIM individuals were still inheriting RNAi.

### Half-life of GFP RNAi memory and statistical analysis

We calculated the GFP RNAi memory half-life values arithmetically based on the raw scoring counts presented in Table S2. For every replicate of very strain in every block, we selected the two scoring time points for which the percentage of GFP-positive individuals were the closest below and above 50, respectively, and calculated the corresponding line equation. Based on this equation, we obtained the theoretical median value of GFP RNAi memory allowing us to generate the boxplots shown in Fig. 1E, Fig. 2B, Fig. 3C, Fig. 4D and Fig. S1 and to conduct statistical analysis. For values oscillating around the median line, a smoothing was realized picking scoring time points further apart. Raw values and indications can be found in Table S3.

Generalized linear mixed model (GLMM) were fit using the glmmTMB (v.1.1.10) package in the R version 4.2.2 (2022-10-31) to compare differences in half-life of GFP RNAi memory across strains or environments.

The model we used to test for the effect of genetic variation was:

model_1 <- glmmTMB(Value ∼ Strain + (1|Block) + (1|Block:Replicate), family = Gamma(link = “log”), data = data)

The model we used to test for the effect of an environmental variable was:

model_2 <- glmmTMB(Value ∼ Strain*Environment + (1|Block) + (1|Block:Replicate), data = data)

We used the emmeans package (v.1.10.6) and Tukey’s post-hoc test for pairwise comparison, either among strains:

emmeans(model, pairwise ∼ Strain, adjust = “tukey”) or between environments for a given strain:

emmeans(model, pairwise ∼ Environment|Strain, adjust = “tukey”)

For blocks realized only once, the random variable (1|Block) was removed from the model. All models were validated by fitting the DHARMa package (v.0.4.7) simulations.

### Haplotype tree building and genetic relatedness

We downloaded the 611 hard-filtered Variant Call Format (VCF) file from CaeNDR (20220216 release) (Cook et al., 2017) from which we extracted the data for our eight strains of interest. We converted this VCF file to a PHILIP file using the vcf2phylip.py script (Ortiz, 2019). The haplotype network was generated using this PHILIP file and the SplitsTree software (v6.4.13) (Huson and Bryant, 2006).

Presence of the drh-1 159 bp deletion among the 611 hard-filtered isotypes was detected using BCFtools (v1.9) and manually checked via CaeNDR’s genome browser tool. Sampling location for each C. elegans isotype was downloaded from CaeNDR (20231213 data release).

## Acknowledgements

We are grateful to Hervé Gendrot for plate pouring and to Aurélien Richaud for laboratory management. We thank Katie Pelletier for help with statistical analysis. We thank Eric Miska, Germano Cecere and members of the Félix lab for discussions. Some strains were provided by the CGC, which is funded by NIH Office of Research Infrastructure Programs (P40 OD010440). We thank the SEGiCel genome editing platform (CNRS, Lyon), especially Arnaud Echard and Margaux Gibert. We thank Wormbase and CaeNDR. This work was supported by grants from the Agence Nationale pour le Recherche ANR-19-CE12-0025-01 and ANR-24-CE12-0362-01. For the purpose of Open Access, the author has applied a CC BY public copyright licence to any Author Accepted Manuscript version arising from this submission.

## Author contributions

M.-A.F. and L.F. designed the project; L.F. performed the introgressions and developed protocols; C. F.-J. provided reagents; M. A. made genome edits for transgene insertion; F.F., M.S. and L.F. performed RNAi memory assays; F.F. performed the statistical analyses and made the figures; F.F., M.-A.F. and M.S. wrote the manuscript with contributions and edits of all authors; M.-A. F. supervised the project, with contributions from L.F. and C. F.-J.

## Supplementary figure legends

**Figure S1: The rank order of memory duration of wild background strains is overall conserved when using the two distinct GFP transgenes**.

(**A**) GFP silencing memory assay comparing in parallel the RNAi memory duration of the reference strain N2 with either the pie-1p::GFP::H2B or the mex-5p::ce-GFP transgenes. E. coli HT115 with the relevant gfp sequence clone for each transgene was fed to initiate RNAi against gfp. Three biological replicates were run for each strain, scoring the proportion of GFP-positive (GFP+) animals at each generation. On the graph, the lines follow the means of three replicates and error bars represent their standard deviation (SD).

(**B**) Boxplot showing the GFP RNAi memory half-lives of a N2 genetic background with the mex-5p::ce-GFP or pie-1p::GFP transgenes. Half-lives were estimated from the scoring data shown in (A).

(**C**) Diagram depicting the two gfp transgenes used in this study. The pie-1 promoter is 2 kb long and not represented to scale. The two transgenes strongly differ in terms of cis-regulatory and coding sequences.

(**D**) Two independent experiments assaying the RNAi memory duration of the same wild isolates containing either the pie-1p::GFP::H2B (Block A) or mex-5p::ce-GFP (Block C, also plotted in Fig. 2) transgenes. For both blocks, E. coli HT115 was fed to initiate RNAi against gfp. As the pie-1p::GFP experiment only tested the MY10, JU1171, JU775, N2 and JU1395 backgrounds, only the relevant strains are plotted for mex-5p::ce-GFP. The variation in memory of the wild strains is overall conserved when using the two distinct GFP transgenes.

(**E**) Boxplots showing the GFP RNAi memory half-lives of the MY10, JU1171, JU775, N2 and JU1395 backgrounds with either GFP transgene. As expected, the memory is longer with the pie-1p::GFP::H2B where piRNA recognition sites have not been avoided.

**Figure S2: Effect of additional culture environments on the duration of RNAi memory**.

(A) Three different experiments (blocks H, I and K) testing for the effect of Chryseobacterium JUb044, Acinetobacter BIGb102, Comamonas BIGb172 and Leucobacter CBX151 on RNAi memory. Here the moderate-memory strain JU1395 or low-memory strain JU1171 containing the mex-5::ce-GFP::tbb-2 transgene were fed with E. coli iOP50 to initiate gfp RNAi. No striking differences were observed in RNAi memory profile between E. coli OP50 or naturally associated bacteria except for a reducing effect of CBX151 on the nematode strain JU1395. Statistics comparing estimated GFP memory half-lives of a given strain in different bacterial condition to the OP50 control: glmm followed by Tukey’s comparison against OP50: p>0.06 for each bacterial strain on JU1171 and p>0.3 for each bacterial strain on JU1395 except with CBX151, p = 0.02. As no major delay was seen in development or egg laying under these conditions, generation time and absolute time are equivalent.

(**B**) The panels correspond to the same block L (distinct from the experiment shown in Figure 4B-C) testing for effect of temperature on RNAi memory in the strains JU1395 and N2 containing the mex-5p::ce-GFP::tbb-2 transgene and fed with E. coli iOP50 to initiate gfp RNAi. Data were plotted either in number of generations or days after RNAi initiation.

(**C**) Independent block J testing for the effect on RNAi memory kinetics of dauer diapause induced at different generations. The long-memory strain XZ1514 containing the mex-5p::ce-GFP::tbb-2 transgene was fed with E. coli iOP50 to initiate gfp RNAi. This experiment was performed at 20°C. Dauer individuals of a given generation were induced by letting parents of the previous generation lay eggs and not re-supplying plates with E. coli OP50. Individuals were left for ∼3 days in the dauer stage. This was performed at every generation by separating in half the control individuals as population to be scored (fed) or population to lay (not fed) (see Methods). Dauer larvae were selected by incubation of the whole population in 1% sodium dodecyl sulfate (SDS) for ∼20 minutes at room temperature with moderate agitation before been deposited back onto standard NGM-OP50 plates to resume development. The dauer diapause does not greatly affect further GFP silencing memory dynamics when expressed in number of generations.

**Figure S3: Sensitivity to RNAi exposure is lower in the C. elegans MY10 strain but the set-24 mutation in the N2 background only affects memory**.

Block F including gfp silencing dynamics during RNAi initiation. Starting from a 15°C stock population where every individual expresses GFP fluorescence, the two-generation (G-1 and G0) of GFP RNAi exposure leads to full GFP silencing except in the MY10 strain. Are shown in the graph the different phases of the gfp silencing inheritance process with initiation of gfp RNAi, establishment of RNAi inheritance, and maintenance of RNAi inheritance.

**Figure S4: Efficiency of gfp RNAi initiation in wild strains and the drh-1 mutant**.

(**A**) Block N showing the dynamics of gfp silencing (generations G-1 and G0) in the MY10, JU1171, N2, drh-1(bab552) in N2, and JU1395 backgrounds harboring the mex-5p::ce-GFP transgene and fed with E. coli iOP50 for RNAi initiation. The drh-1 natural deletion allele reduces both responsiveness to gfp RNAi as well as RNAi memory.

(**B**) Microscopy images illustrating GFP silencing and desilencing dynamics of wild isolates through time. GFP microscopy images from block N taken in parallel with the scoring. Before gfp RNAi, stock populations of all four natural strains show identical patterns of GFP expression. Use of iOP50 with a T444T cassette seems to enhance efficiency of RNAi as even MY10 is almost fully silenced for gfp at G0. At the first generation after RNAi exposure, almost the whole populations of MY10 and JU1171 recover fluorescence while populations of N2 and JU1395 retain some silencing. Dots in the pharynx of silenced individuals are due to autofluorescence.

## Supplementary tables

**Table S1: List of nematode strains used in this paper**.

List of wild C. elegans isolates and their derived transgene-modified counterparts with their origin and method of construction.

**Table S2: Raw scoring data of GFP silencing memory assay**.

Each block corresponds to an independent experiment. DIM and OFF individuals were pooled for simplicity in the graphs. Plasmid code names and bacteria used for GFP RNAi exposure are detailed in Table S4.

**Table S3: Values of GFP RNAi memory half-lives**.

Half-lives of GFP memory were calculated based on the raw scoring counts presented in Table S2 (see methods for procedure). A slash mark in the column for half-lives counted as days corresponds to cases with no difference between the time expressed in generation number or in days. For rare line plots values crossing the 50° value twice, a smoothing of the linear RNAi memory tendency was performed, and indicated by an arrow showing the selected data points.

**Table S4: Bacterial strains and transgenes used in this paper**.

The first sheet shows the list of C. elegans naturally associated-bacteria used as testing environments as well as artificial E. coli strains used for RNAi induction. The second sheet shows the sequences of gfp transgenes and their corresponding targeting dsRNA.

**Table S5: Genotyping of the pie-1p::GFP transgene introgressions**.

As the JU1395 and N2 genetic backgrounds are closely related to each other, we only used primers that could distinguish deletions in the MY10, JU775 and JU1171 isolates compared to N2.

## Notes

### Competing Interest Statement

The authors have declared no competing interest.

## Bibliography

1. *: equal contribution. # correspondence (omitted if last author only)

1. Adrian-Kalchhauser, I., Sultan, S. E., Shama, L. N. S., Spence-Jones, H., Tiso, S., Keller Valsecchi, C. I., et al. (2020). Understanding “non-genetic” inheritance: insights from molecular-evolutionary crosstalk. Trends Ecol. Evol. 35, 1078–1089. doi: 10.1016/j.tree.2020.08.011

2. Ahmed, S.#, and Hodgkin, J. (2000). MRT-2 checkpoint protein is required for germline immortality and telomere replication in C. elegans. Nature 403, 159–164. doi: 10.1038/35003120

3. Aljohani, M. D.*, El Mouridi, S.*, Priyadarshini, M.*, Vargas-Velazquez, A. M.*, and Frøkjær-Jensen, C. (2020). Engineering rules that minimize germline silencing of transgenes in simple extrachromosomal arrays in C. elegans. Nat. Commun. 11, 6300. doi: 10.1038/s41467-020-19898-0

4. Ashe, A.*, Bélicard, T.*, Le Pen, J.*, Sarkies, P.*, Frézal, L., Lehrbach, N. J., et al. (2013). A deletion polymorphism in the Caenorhabditis elegans RIG-I homolog disables viral RNA dicing and antiviral immunity. eLife 2, e00994. doi: 10.7554/eLife.00994

5. Ashe, A., Colot, V., and Oldroyd, B. P. (2021). How does epigenetics influence the course of evolution? Philos. Trans. R. Soc. Lond. B. Biol. Sci. 376, 20200111. doi: 10.1098/rstb.2020.0111

6. Ashe, A.*, Sapetschnig, A.*, Weick, E.-M.*, Mitchell, J.*, Bagijn, M. P., Cording, A. C., et al. (2012). piRNAs can trigger a multigenerational epigenetic memory in the germline of C. elegans. Cell 150, 88–99. doi: 10.1016/j.cell.2012.06.018

7. Batachari, L. E., Dai, A. Y., and Troemel, E. R. (2024). Caenorhabditis elegans RIG-I-like receptor DRH-1 signals via CARDs to activate antiviral immunity in intestinal cells. Proc. Natl. Acad. Sci. 121, e2402126121. doi: 10.1073/pnas.2402126121

8. Baugh, L. R. (2013). To grow or not to grow: nutritional control of development during Caenorhabditis elegans L1 arrest. Genetics 194, 539–555. doi: 10.1534/genetics.113.150847

9. Baugh, L. R.#, and Day, T. (2020). Nongenetic inheritance and multigenerational plasticity in the nematode C. elegans. eLife 9, e58498. doi: 10.7554/eLife.58498

10. Belicard, T., Jareosettasin, P., and Sarkies, P. (2018). The piRNA pathway responds to environmental signals to establish intergenerational adaptation to stress. BMC Biol. 16, 103. doi: 10.1186/s12915-018-0571-y

11. Billi, A. C., Fischer, S. E. J., and Kim, J. K. (2014). Endogenous RNAi pathways in C. elegans. WormBook Online Rev. C. elegans Biol., 1–49. doi: 10.1895/wormbook.1.170.1

12. Bondurianski, R., Day, T. (2018). Extended heredity: a new understanding of inheritance and evolution | Princeton University Press Available at: https://press.princeton.edu/books/hardcover/9780691157672/extended-heredity (Accessed March 10, 2025).

13. Buckley, B. A.*, Burkhart, K. B.*, Gu, S. G., Spracklin, G., Kershner, A., Fritz, H., et al. (2012). A nuclear Argonaute promotes multigenerational epigenetic inheritance and germline immortality. Nature 489, 447–451. doi: 10.1038/nature11352

14. Chen, B., Gilbert, L. A., Cimini, B. A., Schnitzbauer, J., Zhang, W., Li, G.-W., et al. (2013). Dynamic imaging of genomic loci in living human cells by an optimized CRISPR/Cas system. Cell 155, 1479–1491. doi: 10.1016/j.cell.2013.12.001

15. Chen, S., and Phillips, C. M. (2024). HRDE-2 drives small RNA specificity for the nuclear Argonaute protein HRDE-1. Nat. Commun. 15, 957. doi: 10.1038/s41467-024-45245-8

16. Chen, X.*, Wang, K.*, Mufti, F. U. D.*, Xu, D., Zhu, C., Huang, X., et al. (2024). Germ granule compartments coordinate specialized small RNA production. Nat. Commun. 15, 5799. doi: 10.1038/s41467-024-50027-3

17. Chou, H. T., Valencia, F., Alexander, J. C., Bell, A. D., Deb, D., Pollard, D. A., et al. (2024). Diversification of small RNA pathways underlies germline RNA interference incompetence in wild Caenorhabditis elegans strains. Genetics 226, iyad191. doi: 10.1093/genetics/iyad191

18. Coffman, S. R., Lu, J., Guo, X., Zhong, J., Jiang, H., Broitman-Maduro, G., et al. (2017). Caenorhabditis elegans RIG-I homolog mediates antiviral RNA interference downstream of Dicerdependent biogenesis of viral small interfering RNAs. mBio 8, 10.1128/mbio.00264-17.

19. Consalvo, C. D., Aderounmu, A. M.*, Donelick, H. M.*, Aruscavage, P. J., Eckert, D. M., Shen, P. S., et al. (2024). Caenorhabditis elegans Dicer acts with the RIG-I-like helicase DRH-1 and RDE-4 to cleave dsRNA. eLife 13, RP93979. doi: 10.7554/eLife.93979

20. Cook, D. E., Zdraljevic, S., Roberts, J. P., and Andersen, E. C. (2017). CeNDR, the Caenorhabditis elegans natural diversity resource. Nucleic Acids Res. 45, D650–D657. doi: 10.1093/nar/gkw893

21. Crombie, T. A., Zdraljevic, S., Cook, D. E., Tanny, R. E., Brady, S. C., Wang, Y., et al. (2019). Deep sampling of Hawaiian Caenorhabditis elegans reveals high genetic diversity and admixture with global populations. eLife 8, e50465. doi: 10.7554/eLife.50465

22. Cubas, P., Vincent, C., and Coen, E. (1999). An epigenetic mutation responsible for natural variation in floral symmetry. Nature 401, 157–161. doi: 10.1038/43657

23. Day, T.#, and Bonduriansky, R. (2011). A unified approach to the evolutionary consequences of genetic and nongenetic inheritance. Am. Nat. 178, E18–E36. doi: 10.1086/660911

24. de Vanssay, A., Bougé, A.-L., Boivin, A., Hermant, C., Teysset, L., Delmarre, V., et al. (2012). Paramutation in Drosophila linked to emergence of a piRNA-producing locus. Nature 490, 112–115. doi: 10.1038/nature11416

25. Dey, S., Proulx, S. R.#, and Teotónio, H.# (2016). Adaptation to temporally fluctuating environments by the evolution of maternal effects. PLoS Biol. 14, e1002388. doi: 10.1371/journal.pbio.1002388

26. Dirksen, P., Marsh, S. A., Braker, I., Heitland, N., Wagner, S., Nakad, R., et al. (2016). The native microbiome of the nematode Caenorhabditis elegans: gateway to a new host-microbiome model. BMC Biol. 14, 38. doi: 10.1186/s12915-016-0258-1

27. Du, Z.*, Shi, K.*, Brown, J. S., He, T., Wu, W.-S., Zhang, Y., et al. (2023). Condensate cooperativity underlies transgenerational gene silencing. Cell Rep. 42, 112859. doi: 10.1016/j.celrep.2023.112859

28. Duchaine, T. F., Wohlschlegel, J. A., Kennedy, S., Bei, Y., Conte, D., Pang, K., et al. (2006). Functional proteomics reveals the biochemical niche of C. elegans DCR-1 in multiple small-RNA-mediated pathways. Cell 124, 343–354. doi: 10.1016/j.cell.2005.11.036

29. Duempelmann, L.*, Skribbe, M.*, and Bühler, M. (2020). Small RNAs in the transgenerational inheritance of epigenetic information. Trends Genet. 36, 203–214. doi: 10.1016/j.tig.2019.12.001

30. El Mouridi, S., AlHarbi, S., and Frøkjær-Jensen, C. (2021). A histamine-gated channel is an efficient negative selection marker for C. elegans transgenesis. MicroPublication Biol. doi: 10.17912/micropub.biology.000349

31. El Mouridi, S., Alkhaldi, F., and Frøkjær-Jensen, C. (2022). Modular safe-harbor transgene insertion for targeted single-copy and extrachromosomal array integration in Caenorhabditis elegans. G3 GenesGenomesGenetics 12, jkac184. doi: 10.1093/g3journal/jkac184

32. El Mouridi, S., Peng, Y., and Frøkjær-Jensen, C. (2020). Characterizing a strong pan-muscular promoter (Pmlc-1) as a fluorescent co-injection marker to select for single-copy insertions. MicroPublication Biol. doi: 10.17912/micropub.biology.000302

33. Félix, M.-A.#, and Duveau, F. (2012). Population dynamics and habitat sharing of natural populations of Caenorhabditis elegans and C. briggsae. BMC Biol. 10, 59. doi: 10.1186/1741-7007-10-59

34. Fischer, S. E. J.*, Butler, M. D.*, Pan, Q., and Ruvkun, G. (2008). Trans-splicing in C. elegans generates the negative RNAi regulator ERI-6/7. Nature 455, 491–496. doi: 10.1038/nature07274

35. Fischer, S. E. J., Montgomery, T. A., Zhang, C., Fahlgren, N., Breen, P. C., Hwang, A., et al. (2011). The ERI-6/7 helicase acts at the first sage of an siRNA amplification pathway that targets recent gene duplications. PLOS Genet. 7, e1002369. doi: 10.1371/journal.pgen.1002369

36. Fischer, S. E. J.#, and Ruvkun, G.# (2020). Caenorhabditis elegans ADAR editing and the ERI-6/7/MOV10 RNAi pathway silence endogenous viral elements and LTR retrotransposons. Proc. Natl. Acad. Sci. U. S. A. 117, 5987–5996. doi: 10.1073/pnas.1919028117

37. Fitz-James, M. H., and Cavalli, G. (2022). Molecular mechanisms of transgenerational epigenetic inheritance. Nat. Rev. Genet. 23, 325–341. doi: 10.1038/s41576-021-00438-5

38. Frézal, L., Demoinet, E., Braendle, C., Miska, E.#, and Félix, M.-A.# (2018). Natural genetic variation in a multigenerational phenotype in C. elegans. Curr. Biol. 28, 2588–2596.e8. doi: 10.1016/j.cub.2018.05.091

39. Frézal, L.#, and Félix, M.-A.# (2015). C. elegans outside the Petri dish. eLife 4, e05849. doi: 10.7554/eLife.05849

40. Frézal, L.*, Saglio, M.*, Zhang, G., Noble, L., Richaud, A., and Félix, M. (2023). Genome-wide association and environmental suppression of the mortal germline phenotype of wild C. elegans. EMBO Rep. 24, e58116. doi: 10.15252/embr.202358116

41. Frøkjær-Jensen, C., Wayne Davis, M., Hopkins, C. E., Newman, B. J., Thummel, J. M., Olesen, S.-P., et al. (2008). Single-copy insertion of transgenes in Caenorhabditis elegans. Nat. Genet. 40, 1375–1383. doi: 10.1038/ng.248

42. Gibson, D. G., Young, L., Chuang, R.-Y., Venter, J. C., Hutchison, C. A., and Smith, H. O. (2009). Enzymatic assembly of DNA molecules up to several hundred kilobases. Nat. Methods 6, 343–345. doi: 10.1038/nmeth.1318

43. González, R.#, and Félix, M.-A.# (2024). Naturally-associated bacteria modulate Orsay virus infection of Caenorhabditis elegans. PLoS Pathog. 20, e1011947. doi: 10.1371/journal.ppat.1011947

44. Grishok, A., Tabara, H., and Mello, C. C. (2000). Genetic requirements for inheritance of RNAi in C. elegans. Science 287, 2494–2497. doi: 10.1126/science.287.5462.2494

45. Gu, S. G., Pak, J., Guang, S., Maniar, J. M., Kennedy, S., and Fire, A. (2012). Amplification of siRNA in Caenorhabditis elegans generates a transgenerational sequence-targeted histone H3 lysine 9 methylation footprint. Nat. Genet. 44, 157–164. doi: 10.1038/ng.1039

46. Heard, E.#, and Martienssen, R. A.# (2014). Transgenerational epigenetic inheritance: myths and mechanisms. Cell 157, 95–109. doi: 10.1016/j.cell.2014.02.045

47. Helanterä, H.#, and Uller, T. (2010). The Price Equation and Extended Inheritance. Philos. Theory Biol. 2. doi: 10.3998/ptb.6959004.0002.001

48. Hibshman, J. D*., Webster, A. K.*, and Baugh, L. R. (2021). Liquid-culture protocols for synchronous starvation, growth, dauer formation, and dietary restriction of Caenorhabditis elegans. STAR Protoc. 2, 100276. doi: 10.1016/j.xpro.2020.100276

49. Hodgkin, J.#, Félix, M.-A., Clark, L. C., Stroud, D., and Gravato-Nobre, M. J. (2013). Two Leucobacter strains exert complementary virulence on Caenorhabditis including death by worm-star formation. Curr. Biol. 23, 2157–2161. doi: 10.1016/j.cub.2013.08.060

50. Houri-Zeevi, L.#, Korem Kohanim, Y., Antonova, O., and Rechavi, O.# (2020). Three rules explain transgenerational small RNA inheritance in C. elegans. Cell 182, 1186–1197.e12. doi: 10.1016/j.cell.2020.07.022

51. Houri-Ze’evi, L., Korem, Y., Sheftel, H., Faigenbloom, L., Toker, I. A., Dagan, Y., et al. (2016). A tunable mechanism determines the duration of the transgenerational small RNA inheritance in C. elegans. Cell 165, 88–99. doi: 10.1016/j.cell.2016.02.057

52. Houri-Zeevi, L.#, Teichman, G., Gingold, H., and Rechavi, O.# (2021). Stress resets ancestral heritable small RNA responses. eLife 10, e65797. doi: 10.7554/eLife.65797

53. Hu, P. J. (2007). Dauer. WormBook Online Rev. C elegans Biol., 1–19. doi: 10.1895/wormbook.1.144.1

54. Huson, D. H., and Bryant, D. (2006). Application of phylogenetic networks in evolutionary studies. Mol. Biol. Evol. 23, 254–267. doi: 10.1093/molbev/msj030

55. Jablonka, E., Oborny, B., Molnár, I., Kisdi, E., Hofbauer, J., and Czárán, T. (1995). The adaptive advantage of phenotypic memory in changing environments. Philos. Trans. R. Soc. Lond. B. Biol. Sci. 350, 133–141. doi: 10.1098/rstb.1995.0147

56. Karin, O.#, Miska, E. A., and Simons, B. D.# (2023). Epigenetic inheritance of gene silencing is maintained by a self-tuning mechanism based on resource competition. Cell Syst. 14, 24–40.e11. doi: 10.1016/j.cels.2022.12.003

57. Karp, X. (2018). Working with dauer larvae. WormBook Online Rev. C. elegans Biol. 2018, 1–19. doi: 10.1895/wormbook.1.180.1

58. Kennedy, S., Wang, D., and Ruvkun, G. (2004). A conserved siRNA-degrading RNase negatively regulates RNA interference in C. elegans. Nature 427, 645–649. doi: 10.1038/nature02302

59. Klosin, A., Casas, E., Hidalgo-Carcedo, C., Vavouri, T.#, and Lehner, B.# (2017). Transgenerational transmission of environmental information in C. elegans. Science 356, 320–323. doi: 10.1126/science.aah6412

60. Knudsen-Palmer, D. R., Raman, P., Ettefa, F., De Ravin, L., and Jose, A. M. (2024). Target-specific requirements for RNA interference can arise through restricted RNA amplification despite the lack of specialized pathways. eLife 13, RP97487. doi: 10.7554/eLife.97487

61. Lachmann, M., and Jablonka, E. (1996). The inheritance of phenotypes: an adaptation to fluctuating environments. J. Theor. Biol. 181, 1–9. doi: 10.1006/jtbi.1996.0109

62. Lee, R. C., Hammell, C. M., and Ambros, V. (2006). Interacting endogenous and exogenous RNAi pathways in Caenorhabditis elegans. RNA 12, 589–597. doi: 10.1261/rna.2231506

63. Lev, I.*, Seroussi, U.*, Gingold, H., Bril, R., Anava, S., and Rechavi, O. (2017). MET-2-dependent H3K9 methylation suppresses transgenerational small RNA inheritance. Curr. Biol. 27, 1138–1147. doi: 10.1016/j.cub.2017.03.008

64. Lind, M. I.*#, and Spagopoulou, F.*# (2018). Evolutionary consequences of epigenetic inheritance. Heredity 121, 205–209. doi: 10.1038/s41437-018-0113-y

65. Mao, K., Breen, P., and Ruvkun, G. (2020). Mitochondrial dysfunction induces RNA interference in C. elegans through a pathway homologous to the mammalian RIG-I antiviral response. PLOS Biol. 18, e3000996. doi: 10.1371/journal.pbio.3000996

66. Mello, C. C., Kramer, J. M., Stinchcomb, D., and Ambros, V. (1991). Efficient gene transfer in C. elegans: extrachromosomal maintenance and integration of transforming sequences. EMBO J. 10, 3959–3970. doi: 10.1002/j.1460-2075.1991.tb04966.x

67. Merritt, C., Rasoloson, D., Ko, D., and Seydoux, G. (2008). 31 UTRs are the primary regulators of gene expression in the C. elegans germline. Curr. Biol. 18, 1476–1482. doi: 10.1016/j.cub.2008.08.013

68. Miska, E. A., and Ferguson-Smith, A. C. (2016). Transgenerational inheritance: models and mechanisms of non-DNA sequence-based inheritance. Science 354, 59–63. doi: 10.1126/science.aaf4945

69. Ortiz, E. M. (2019). vcf2phylip v2.0: convert a VCF matrix into several matrix formats for phylogenetic analysis. doi: 10.5281/zenodo.2540861

70. Ouyang, J. P. T., Zhang, W. L., and Seydoux, G. (2022). The conserved helicase ZNFX-1 memorializes silenced RNAs in perinuclear condensates. Nat. Cell Biol. 24, 1129– 1140. doi: 10.1038/s41556-022-00940-w

71. Paaby, A. B.#, White, A. G., Riccardi, D. D., Gunsalus, K. C., Piano, F., and Rockman, M. V.# (2015). Wild worm embryogenesis harbors ubiquitous polygenic modifier variation. eLife 4, e09178. doi: 10.7554/eLife.09178

72. Pak, J., and Fire, A. (2007). Distinct Populations of Primary and Secondary Effectors During RNAi in C. elegans. Science 315, 241–244. doi: 10.1126/science.1132839

73. Pak, J., Maniar, J. M., Mello, C. C., and Fire, A. (2012). Protection from Feed-Forward Amplification in an Amplified RNAi Mechanism. Cell 151, 885–899. doi: 10.1016/j.cell.2012.10.022

74. Pollard, D. A., and Rockman, M. V. (2013). Resistance to germline RNA interference in a Caenorhabditis elegans wild isolate exhibits complexity and nonadditivity. G3 Bethesda Md 3, 941–947. doi: 10.1534/g3.113.005785

75. Radman, I., Greiss, S.#, and Chin, J. W.# (2013). Efficient and rapid C. elegans transgenesis by bombardment and hygromycin B selection. PLOS ONE 8, e76019. doi: 10.1371/journal.pone.0076019

76. Rivoire, O., and Leibler, S. (2014). A model for the generation and transmission of variations in evolution. Proc. Natl. Acad. Sci. U. S. A. 111, E1940–1949. doi: 10.1073/pnas.1323901111

77. Samuel, B. S., Rowedder, H., Braendle, C., Félix, M.-A.#, and Ruvkun, G.# (2016). Caenorhabditis elegans responses to bacteria from its natural habitats. Proc. Natl. Acad. Sci. 113, E3941–E3949. doi: 10.1073/pnas.1607183113

78. Schott, D.#, Yanai, I., and Hunter, C. P.# (2014). Natural RNA interference directs a heritable response to the environment. Sci. Rep. 4, 7387. doi: 10.1038/srep07387

79. Schulenburg, H.#, and Félix, M.-A.# (2017). The natural biotic environment of Caenorhabditis elegans. Genetics 206, 55–86. doi: 10.1534/genetics.116.195511

80. Shukla, A., Perales, R., and Kennedy, S. (2021). piRNAs coordinate poly(UG) tailing to prevent aberrant and perpetual gene silencing. Curr. Biol. 31, 4473–4485.e3. doi: 10.1016/j.cub.2021.07.076

81. Sijen, T.*, Fleenor, J.*, Simmer, F.*, Thijssen, K. L., Parrish, S., Timmons, L., et al. (2001). On the Role of RNA Amplification in dsRNA-Triggered Gene Silencing. Cell 107, 465–476. doi: 10.1016/S0092-8674(01)00576-1

82. Sowa, J. N., Jiang, H., Somasundaram, L., Tecle, E., Xu, G., Wang, D., et al. (2020). The Caenorhabditis elegans RIG-I homolog DRH-1 mediates the intracellular pathogen response upon viral infection. J. Virol. 94, 10.1128/jvi.01173-19.

83. Spracklin, G., Fields, B., Wan, G., Becker, D., Wallig, A., Shukla, A., et al. (2017). The RNAi inheritance machinery of Caenorhabditis elegans. Genetics 206, 1403–1416. doi: 10.1534/genetics.116.198812

84. Stiernagle, T. (2006). Maintenance of C. elegans. WormBook Online Rev. C elegans Biol., 1–11. doi: 10.1895/wormbook.1.101.1

85. Sturm, Á.*, Saskői, É.*, Tibor, K., Weinhardt, N., and Vellai, T. (2018). Highly efficient RNAi and Cas9-based auto-cloning systems for C. elegans research. Nucleic Acids Res. 46, e105. doi: 10.1093/nar/gky516

86. Tabara, H., Yigit, E., Siomi, H., and Mello, C. C. (2002). The dsRNA binding protein RDE-4 interacts with RDE-1, DCR-1, and a DExH-Box Helicase to Direct RNAi in C. elegans. Cell 109, 861–871. doi: 10.1016/S0092-8674(02)00793-6

87. Timmons, L., Court, D. L., and Fire, A. (2001). Ingestion of bacterially expressed dsRNAs can produce specific and potent genetic interference in Caenorhabditis elegans. Gene 263, 103–112. doi: 10.1016/S0378-1119(00)00579-5

88. Tsai, H.-Y., Chen, C.-C. G., Conte, D., Moresco, J. J., Chaves, D. A., Mitani, S., et al. (2015). A Ribonuclease Coordinates siRNA Amplification and mRNA Cleavage during RNAi. Cell 160, 407–419. doi: 10.1016/j.cell.2015.01.010

89. Vasale, J. J.*, Gu, W.*, Thivierge, C., Batista, P. J., Claycomb, J. M., Youngman, E. M., et al. (2010). Sequential rounds of RNA-dependent RNA transcription drive endogenous small-RNA biogenesis in the ERGO-1/Argonaute pathway. Proc. Natl. Acad. Sci. 107, 3582–3587. doi: 10.1073/pnas.0911908107

90. Vastenhouw, N. L., Brunschwig, K., Okihara, K. L., Müller, F., Tijsterman, M., and Plasterk, R. H. A. (2006). Gene expression: long-term gene silencing by RNAi. Nature 442, 882. doi: 10.1038/442882a

91. Weigel, D.#, and Colot, V.# (2012). Epialleles in plant evolution. Genome Biol. 13, 249. doi: 10.1186/gb-2012-13-10-249

92. Woodhouse, R. M., Buchmann, G., Hoe, M., Harney, D. J., Low, J. K. K., Larance, M., et al. (2018). Chromatin modifiers SET-25 and SET-32 are required for establishment but not long-term maintenance of transgenerational epigenetic inheritance. Cell Rep. 25, 2259–2272.e5. doi: 10.1016/j.celrep.2018.10.085

93. Wu, X., Shi, Z., Cui, M., Han, M., and Ruvkun, G. (2012). Repression of Germline RNAi Pathways in Somatic Cells by Retinoblastoma Pathway Chromatin Complexes. PLOS Genet. 8, e1002542. doi: 10.1371/journal.pgen.1002542

94. Xu, F.*, Feng, X.*, Chen, X., Weng, C., Yan, Q., Xu, T., et al. (2018). A Cytoplasmic Argonaute Protein Promotes the Inheritance of RNAi. Cell Rep. 23, 2482–2494. doi: 10.1016/j.celrep.2018.04.072

95. Zeiser, E., Frøkjær-Jensen, C., Jorgensen, E., and Ahringer, J. (2011). MosSCI and Gateway compatible plasmid toolkit for constitutive and inducible expression of transgenes in the C. elegans germline. PLOS ONE 6, e20082. doi: 10.1371/journal.pone.0020082

96. Zeng, C.*, Furlan, G.*, Almeida, M. V.*, Rueda-Silva, J. C., Price, J., Mars, J., et al. (2025). A SET domain-containing protein and HCF-1 maintain transgenerational epigenetic memory. 2025.03.25.645221. doi: 10.1101/2025.03.25.645221

97. Zhang, G.#, Félix, M.-A.#, and Andersen, E. C.# (2024). Transposon-mediated genic rearrangements underlie variation in small RNA pathways. Sci. Adv. 10, eado9461. doi: 10.1126/sciadv.ado9461

