## Supplementary figures and images for "Intraspecific variation in the duration of epigenetic inheritance"

### Supplementary Figure 1

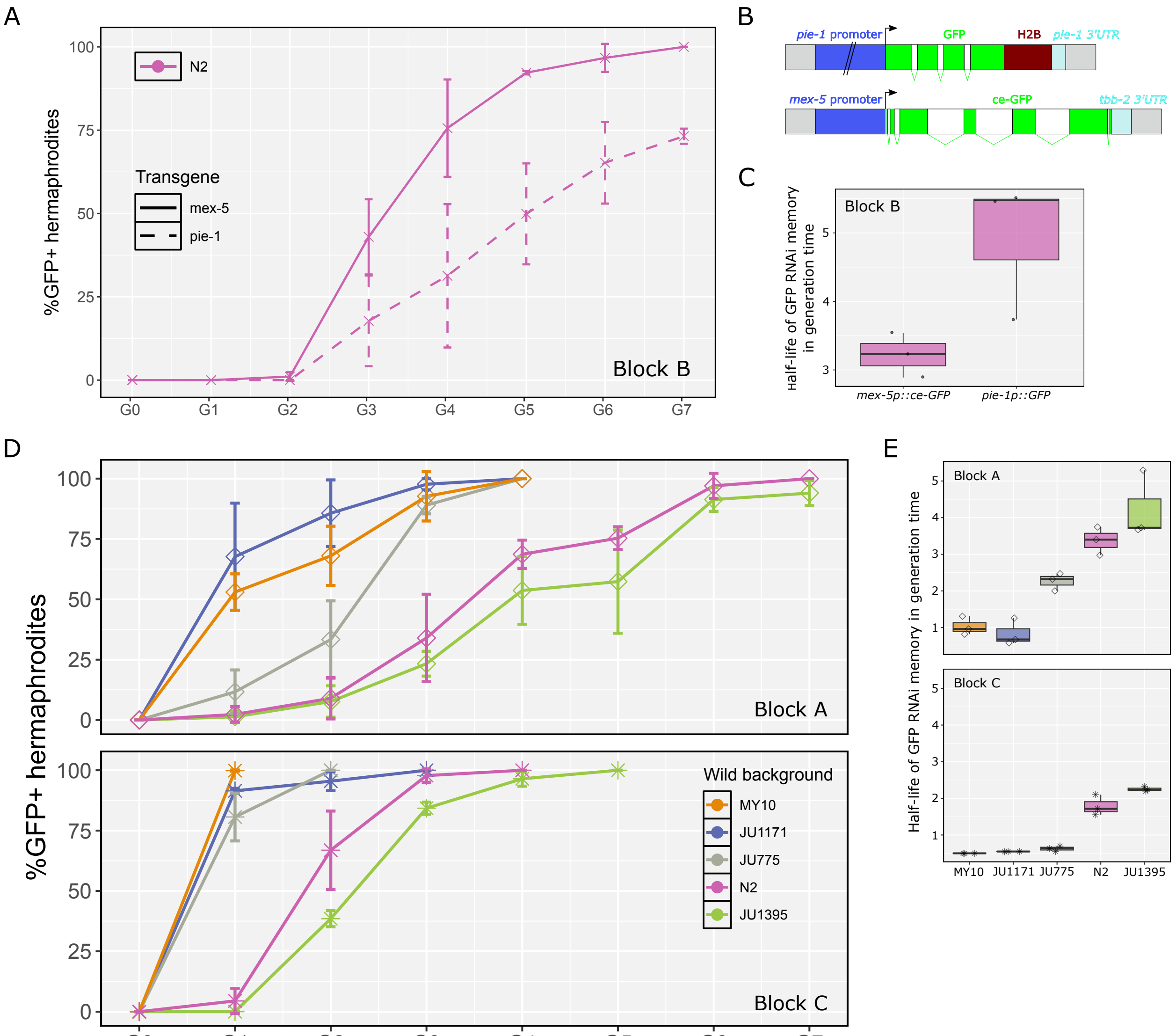

### Supplementary Figure 2

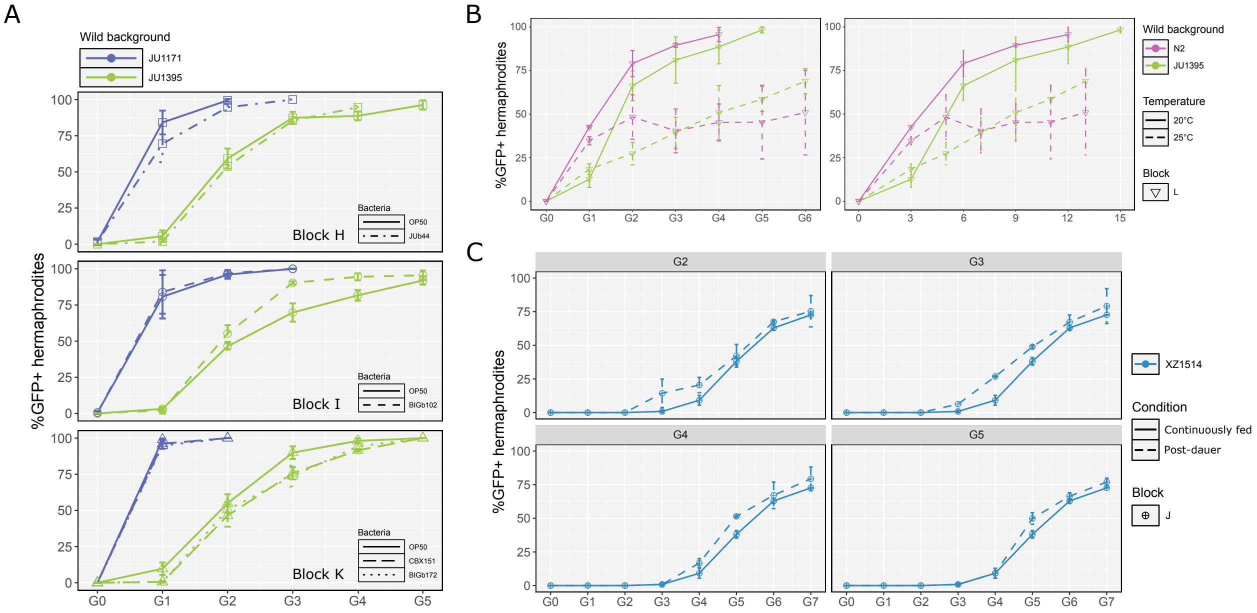

### Supplementary Figure 3

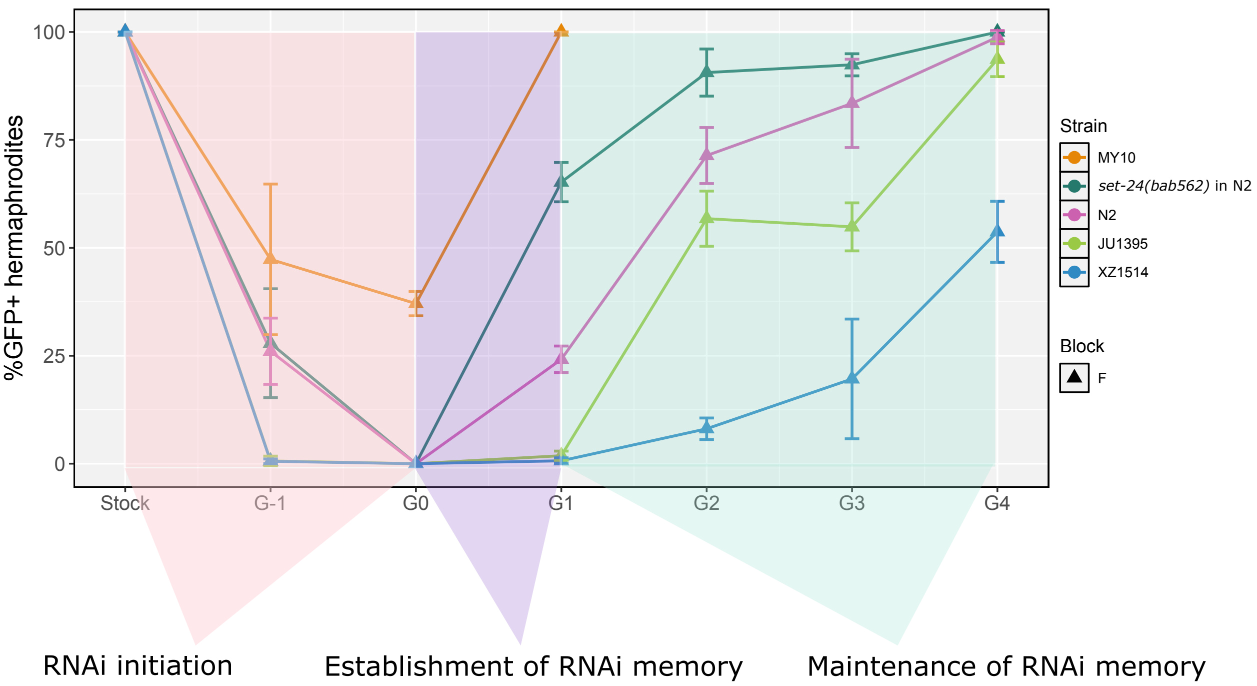

### Supplementary Figure 4

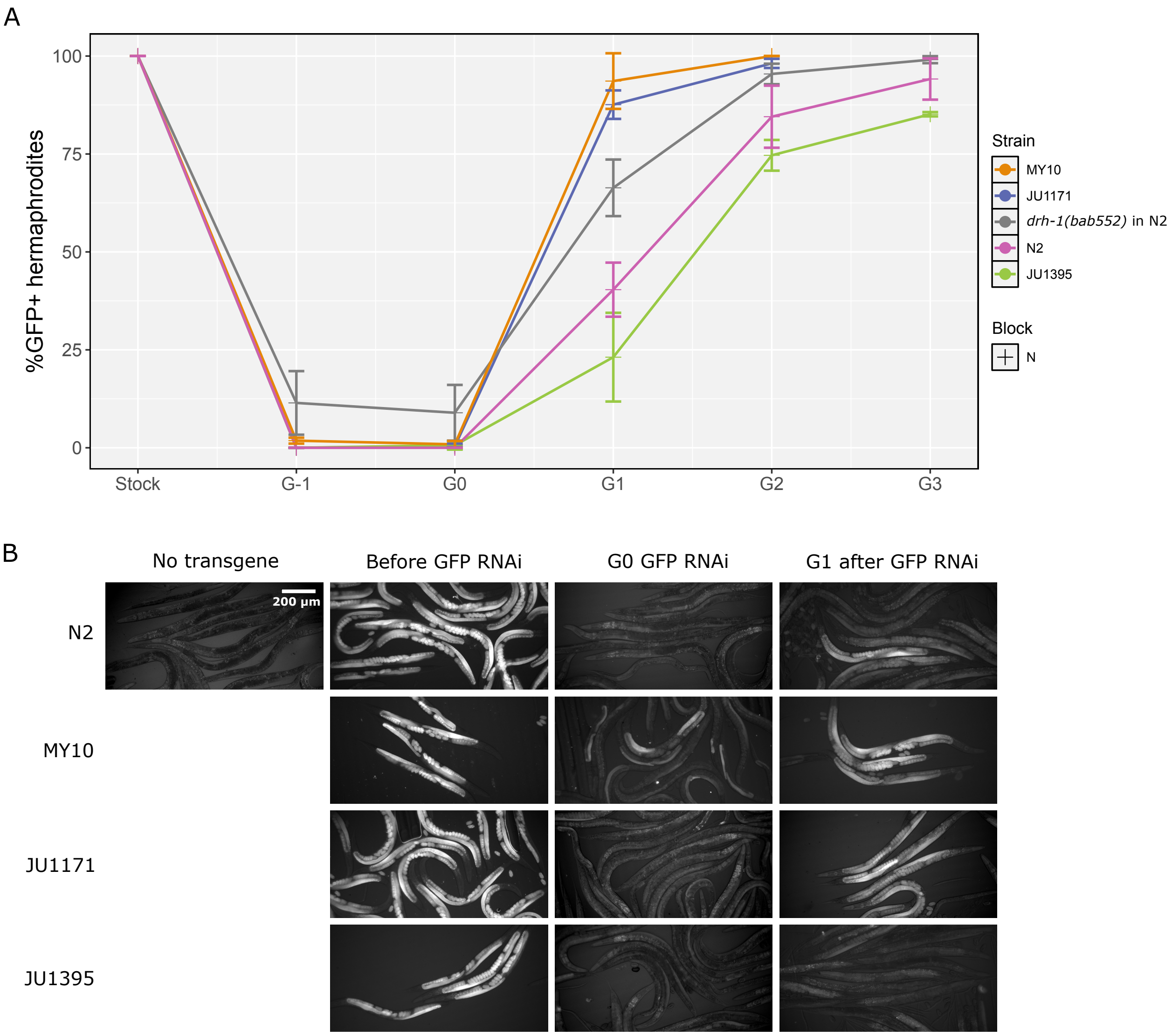
